# Genetic regulation of ALK inhibitor response in diverse human cell types

**DOI:** 10.64898/2026.11.16.766986

**Authors:** Elizabeth Geena Woo, Katherine L. Rhodes, Natalia M. Gonzales, Kenneth A. Barr, Emaan Mohsin, Everett E. Vokes, Yoav Gilad

## Abstract

Anaplastic lymphoma kinase (ALK) inhibitors are effective treatments for ALK-positive non-small cell lung cancer but cause diverse adverse events affecting multiple tissues. Genetic variation may contribute to inter-individual differences in susceptibility, but the small number of treated patients limits the power of genetic association studies. Moreover, the diversity of affected tissues and cell types makes functional investigation challenging. We used induced pluripotent stem cells from 49 individuals to generate heterogeneous differentiating cultures (HDCs) containing multiple cell types, treated them with five ALK inhibitors (crizotinib, alectinib, ceritinib, brigatinib, and lorlatinib) for 24 hours, and characterized transcriptional responses using single-cell RNA sequencing. These responses showed cell-type-specific and shared transcriptional effects. Integrating gene expression with individual genotypes, we identified 4,371 expression quantitative trait loci (eQTLs) across 21 cell types, including 2,137 response eQTLs whose regulatory effects differ between untreated and drug-treated cells. These results nominate candidates for future studies of ALK inhibitor toxicity.

## Introduction

Conventional chemotherapeutic drugs are designed to kill tumors by interfering with metabolism, inhibiting mitosis, and promoting cell death^1^. Due to their broad, nonspecific molecular effects, these drugs also tend to harm healthy tissues. For example, the anthracycline doxorubicin, which induces oxidative stress and interferes with DNA synthesis, replication, and transcription^2^, is notorious for adverse cardiovascular events that often result in poor treatment adherence and reduced efficacy^3^. In contrast, targeted therapies are designed to inhibit specific oncogenic malignancies expressed in tumor cells rather than broadly disrupting processes shared by all dividing cells^4^. For example, ∼5% of non-small cell lung cancer (NSCLC) patients have tumors that harbor genomic rearrangements involving anaplastic lymphoma kinase (*ALK*)^5,6^, most commonly an *EML4*-*ALK* fusion.^7^ This rearrangement leads to constitutive ALK activity and overactivation of downstream pathways involved in cell growth, survival, and metabolism^8^, ultimately resulting in tumors. While ALK inhibitors are more effective than conventional drugs for treating ALK-positive lung cancer^9^, they still produce unwanted side effects in many patients, some of which are severe enough to suspend treatment^10^.

The ability to predict a patient’s adverse response to specific drugs would allow physicians to develop safer and more effective treatment plans. Sex, age, medical history, lifestyle, and environmental factors can all affect how an individual responds to a given drug. These responses may also be modulated by genetic variants, which hold great potential for the future of precision medicine^11,12^. The problem is that these variants are difficult to identify. Genome-wide association studies (GWAS) provide an unbiased way to identify loci associated with drug effects, but they also require large patient samples that are impossible to obtain for ALK inhibitors and other drugs designed to target specific or rare cancer subtypes. An alternative is to search for loci that interact with drug treatment (genotype-by-drug interactions). Such loci may directly or indirectly influence an individual’s response to treatment and therefore represent promising candidates for predicting drug-induced adverse effects. One way to identify these interactions is through response expression quantitative trait locus (response eQTL) mapping, which tests for genetic variants whose association with gene expression changes following drug treatment^13^. By mapping response eQTLs across cell types relevant to drug-induced side effects we can identify genes and pathways that contribute to variation in drug responses and nominate candidate biomarkers for predicting clinical outcomes in the general population.

In previous work, Knowles *et al.* identified hundreds of doxorubicin response eQTLs using a panel of differentiated human cardiomyocytes^14^. In addition to predicting signatures of *in vitro* cardiotoxicity, we used these eQTLs to retrospectively predict doxorubicin-induced cardiotoxicity *in vivo*. Here, we extend the premise established by Knowles *et al.* to characterize the *in vitro* responses of human cells to five ALK-inhibiting drugs: crizotinib, ceritinib, alectinib, brigatinib, and lorlatinib. ALK is expressed in the developing and adult central nervous system (CNS), and although it has a known role in neural development, its physiological function in adult tissues is not well-characterized^15,16^. ALK inhibitors can affect normal activity of ALK in non-cancer cells and act on other kinases in addition to ALK (off-target effects), but how these effects modulate individual- and tissue-specific risk for adverse events is unclear^17^. ALK inhibitors also differ in chemical structure, off-target kinase activity, and pharmacokinetic properties, which could account for their differing toxicity profiles (Table 1).

**Table 1.** Adverse events listed in FDA information sheets Section 5 (Warnings and Precautions) for each ALK inhibitor. Asterisks (*) denote the most common adverse reactions (at least 20-25%) in adult patients with NSCLC as noted in Section 6 (Adverse Reactions), excluding general systemic effects such as edema, fatigue, cough, etc. Incidence is from reported numbers in Section 5 (Warnings and Precautions) and Section 6 (Adverse Reactions), primarily based on pooled safety population or clinical trial data. Cell types listed are those found in HDCs that are plausibly relevant to tissues affected by the corresponding adverse events, and cell types in which we mapped eQTLs are bolded. Grades refer to Common Terminology Criteria for Adverse Events (CTCAE), where Grade 1 is mild, Grade 2 is moderate, Grade 3 is severe/medically significant, Grade 4 is life-threatening requiring urgent intervention, and Grade 5 results in death related to the adverse event.

| Drug (generation) | Additional non-ALK targets | Adverse events | Reported incidence | HDC cell types plausibly relevant to affected tissue |
|---|---|---|---|---|
| Crizotinib (1<sup>st</sup> gen) | MET, ROS1 | Hepatotoxicity (elevated AST/ALT)* | 11% (ALT >5x upper limit of normal)<br>6% (AST >5x upper limit of normal) | **Hepatoblasts**<br>Stellate cells<br>Ductal cells |
|  |  | Interstitial lung disease/pneumonitis | Any grade: 2.9%<br>Grade 3 or 4: 1%<br>Fatal: 0.5% | **Bronchiolar/alveolar epithelial cells**<br>**Ciliated epithelial cells**<br>**Stromal cells**<br>Myeloid cells |
|  |  | QT interval prolongation | All-grade: 2.1% | **Cardiac mesoderm cells**<br>Cardiomyocytes<br>Endocardial cells |
|  |  | Bradycardia | All-grade: 13%<br>Grade 3: 2.4% | **Cardiac mesoderm cells**<br>Cardiomyocytes<br>Endocardial cells |
|  |  | Vision disorder (diplopia, photophobia, photopsia, reduced visual acuity, blurred vision, vitreous floaters, visual impairment)* | All-grade: 60-71% (in FDA info sheet Section 6-reported studies)<br>Grade 4: 0.2% | **Retinal pigment cells**<br>**Retinal neurons**<br>**Retinal progenitors/Müller glia**<br>Lens fibre cells |
|  |  | Gastrointestinal (GI) toxicity (nausea, diarrhea, vomiting, constipation)*, particularly for pediatric/young adult patients with anaplastic large cell lymphoma (ALCL) and inflammatory myofibroblastic tumors (IMT) | All-grade: >60% (in FDA Section 6-reported studies, not limited to just ALCL or IMT patients) |  |
| Alectinib (2<sup>nd</sup> gen) | RET | Hepatotoxicity* | All-grade: 41%<br>Grade 3+: 8% | **Hepatoblasts**<br>Stellate cells<br>Ductal cells |
|  |  | Interstitial lung disease/pneumonitis | All-grade: 1.3%<br>Grade 3+ : 0.4% | **Bronchiolar/alveolar epithelial cells**<br>**Ciliated epithelial cells**<br>**Stromal cells**<br>Myeloid cells |
|  |  | Renal impairment | All-grade: 12%<br>Grade 3+: 1.7% (0.4% fatal) | Metanephric cells<br>Stromal cells |
|  |  | Bradycardia | All-grade: 11% | **Cardiac mesoderm cells**<br>Cardiomyocytes<br>Endocardial cells |
|  |  | Myalgia* | All-grade: 31%<br>Grade 3+: 0.8% | Skeletal muscle cells<br>Satellite cells |
|  |  | Creatine phosphokinase (CPK) elevation | All-grade: 56%<br>Grade 3+: 6% | Skeletal muscle cells<br>Satellite cells |
|  |  | Hemolytic anemia | All-grade: 3.1% | Erythroblasts |
| Brigatinib (2<sup>nd</sup> gen) | ROS1, IGF-1R, FLT3, EGFR | Interstitial lung disease/pneumonitis | All-grade: 5.1%<br>Grade 3+: 2.2% | **Bronchiolar/alveolar epithelial cells**<br>**Ciliated epithelial cells**<br>**Stromal cells**<br>Myeloid cells |
|  |  | Hypertension* | All-grade: 32%<br>Grade 3+: 13% | **Vascular endothelial cells** |
|  |  | Bradycardia | All grade: 8.1%<br>Grade 3+: 0.7% | **Cardiac mesoderm cells**<br>Cardiomyocytes<br>Endocardial cells |
|  |  | Visual disturbance (blurred vision, photophobia, photopsia, reduced visual acuity) | Grade 1 or 2: 7.4% | **Retinal pigment cells**<br>**Retinal neurons**<br>**Retinal progenitors/Müller glia**<br>Lens fibre cells |
|  |  | Creatine phosphokinase (CPK) elevation | All-grade: 81%<br>Grade 3+: 24% | Skeletal muscle cells<br>Satellite cells |
|  |  | Pancreatic enzymes elevation (lipase/amylase) | All-grade: 52% (amylase), 59% (lipase)<br>Grade 3+ 6.8% (amylase), 17% (lipase) | Acinar cells<br>Ductal cells |
|  |  | Hepatotoxicity | All-grade: 72% (AST elevation), 52% (ALT elevation)<br>Grade 3+: 4.5% (AST elevation), 5.2% (ALT elevation) | **Hepatoblasts**<br>Stellate cells<br>Ductal cells |
|  |  | Hyperglycemia | All-grade: 56%<br>Grade 3+: 7.5% | **Hepatoblasts**<br>**Epicardial fat cells**<br>Islet endocrine cells |
|  |  | Photosensitivity/rash* | All-grade: 3.7%<br>Grade 3+: 0.7% | **Squamous epithelial cells** |
| Ceritinib (2<sup>nd</sup> gen) | IGF-1R, ROS1, STK22D | Gastrointestinal (nausea, vomiting, diarrhea)* | All-grade: 95%<br>Grade 3+: 14% | **ENS glia**<br>**Goblet cells** |
|  |  | Hepatotoxicity | All-grade: 16% | **Hepatoblasts**<br>Stellate cells<br>Ductal cells |
|  |  | Interstitial lung disease/pneumonitis | All-grade: 2.4%<br>Grade 3+: 1.3% | **Bronchiolar/alveolar epithelial cells**<br>**Ciliated epithelial cells**<br>**Stromal cells**<br>Myeloid cells |
|  |  | QT interval prolongation | All-grade: 6% | **Cardiac mesoderm cells**<br>Cardiomyocytes<br>Endocardial cells |
|  |  | Hyperglycemia | Grade 3+: 13% | **Hepatoblasts**<br>**Epicardial fat cells**<br>Islet endocrine cells |
|  |  | Bradycardia | All-grade: 1.1% | **Cardiac mesoderm cells**<br>Cardiomyocytes<br>Endocardial cells |
|  |  | Pancreatic enzyme elevation (lipase/amylase) | Grade 3+: 7% (amylase), 14% (lipase) | Acinar cells<br>Ductal cells |
| Lorlatinib (3<sup>rd</sup> gen) | ROS1 | Hepatotoxicity (AST/ALT elevations) | All-grade: 48% (AST), 37% (ALT)<br>Grade 3+: 2.7% (AST), 0.7% (ALT) | **Hepatoblasts**<br>Stellate cells<br>Ductal cells |
|  |  | CNS effects*<br>Cognitive effects<br>Mood effects<br>Speech effects<br>Psychotic effects<br>Mental status changes<br>Seizures<br>Sleep effects | All-grade: 52%<br>28% (2.9% severe)<br>21% (1.7% severe)<br>11% (0.6% severe)<br>7% (0.6% severe)<br>1.3%(1.1% severe)<br>1.9%<br>12% | **Radial glia**<br>**Neuronal IPC**<br>**Neuroblast**<br>**Neuron**<br>**Glioblast**<br>Oligodendrocytes |
|  |  | Hyperlipidemia* | Grade 3+: 18% (total cholesterol), 19% (triglycerides) | **Hepatoblasts**<br>**Epicardial fat cells**<br>**Vascular endothelial cells**<br>Smooth cells<br>Myeloid cells |
|  |  | Atrioventricular block | All-grade: 1.9%<br>Grade 3+: 0.2% | **Cardiac mesoderm cells**<br>Cardiomyocytes<br>Endocardial cells |
|  |  | Interstitial lung disease/pneumonitis | All-grade: 1.9%<br>Grade 3+: 0.6% | **Bronchiolar/alveolar epithelial cells**<br>**Ciliated epithelial cells**<br>**Stromal cells**<br>Myeloid cells |
|  |  | Hypertension | All-grade: 13%<br>Grade 3+: 6% | **Vascular endothelial cells**<br>Smooth muscle cells<br>Metanephric cells |
|  |  | Hyperglycemia | All-grade: 9%<br>Grade 3+: 3.2% | **Hepatoblasts**<br>**Epicardial fat cells**<br>Islet endocrine cells |

While ALK inhibitors share many adverse events as a class, they differ in the relative prominence of some adverse events (Table 2). For example, crizotinib, a first-generation ALK inhibitor originally developed against *MET* and also active against *ROS1,* often produces visual disturbances^18^. Second-generation inhibitors (ceritinib, alectinib, and brigatinib) were developed to overcome crizotinib resistance and improve delivery to the central nervous system, which is important because the brain is a common site of metastases^19–21^. Among these, ceritinib is characteristically associated with gastrointestinal events^22^, alectinib with hepatic effects and myalgia^23^, and brigatinib with early-onset pulmonary events and hypertension^24^. Lorlatinib, a third-generation ALK/ROS1 inhibitor designed for high blood-brain penetration and activity against ALK resistance mutations, is distinctively associated with hyperlipidemia and neurocognitive and mood effects^25–27^. It is not well-understood how each drug produces its particular pattern of tissue-specific toxicity and why susceptibility differs between individuals. Transcriptional responses to ALK inhibition have been studied in cancer cells, mostly in the context of drug resistance^28–30^, but responses in non-cancer cell types relevant to these adverse events remain largely unexplored.

**Table 2.**
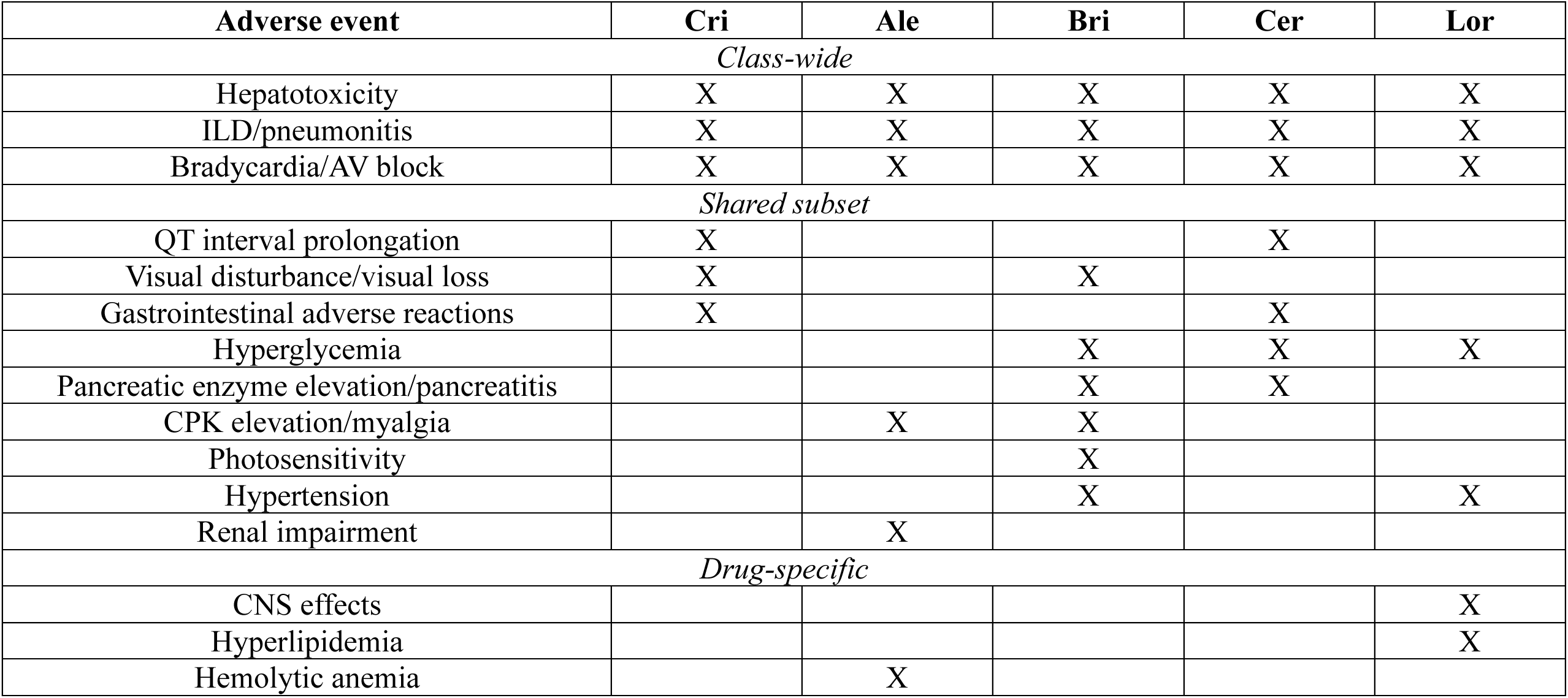
Class- and drug-specific adverse events. X indicates the adverse event is a warning in Section 5 of the FDA label. This Supplementary Data summarizes the clinically actionable dose modification events listed in the Warnings and Precautions section, and is therefore not a comprehensive list of all adverse events that have been reported for each drug.

To capture a comprehensive spectrum of genetic risk for adverse events induced by ALK inhibitors, we needed an *in vitro* system capable of modeling drug-induced perturbations across multiple cell types. Using a panel of 49 human induced pluripotent stem cell lines (iPSCs), we generated heterogeneous differentiating cultures (HDCs), which consist of cells from each of the three germ layers, including cell types representative of tissues affected by ALK inhibitor toxicity^31,32^ (Table 1). To characterize inter-individual transcriptional responses to ALK inhibitors, we independently treated HDCs with each of the five ALK inhibitors and used single-cell RNA-sequencing (scRNA-seq) to capture their transcriptional profiles after 24 hours of drug exposure.

## Results

Using the unguided differentiation approach described by Popp *et al.,* we generated HDCs from an existing panel of 49 human iPSCs^33^. After culturing HDCs for 20 days, we pooled cell lines within each batch in equal proportions, divided them into five tubes, and administered a predetermined dose of crizotinib, ceritinib, alectinib, brigatinib, or lorlatinib to each tube (Fig. 1a, see Methods for a description of our pilot study). Because our primary goal was to identify *cis*-eQTLs within each treatment, we elected to use existing scRNA-seq data from untreated, donor-matched HDCs^32^ as the control group for our eQTL analysis (i.e., rather than collecting additional data from untreated HDCs). Our previous work has shown that baseline eQTLs identified in HDCs are robust with respect to collection batch and highly reproducible^34^.

**Figure 1.**
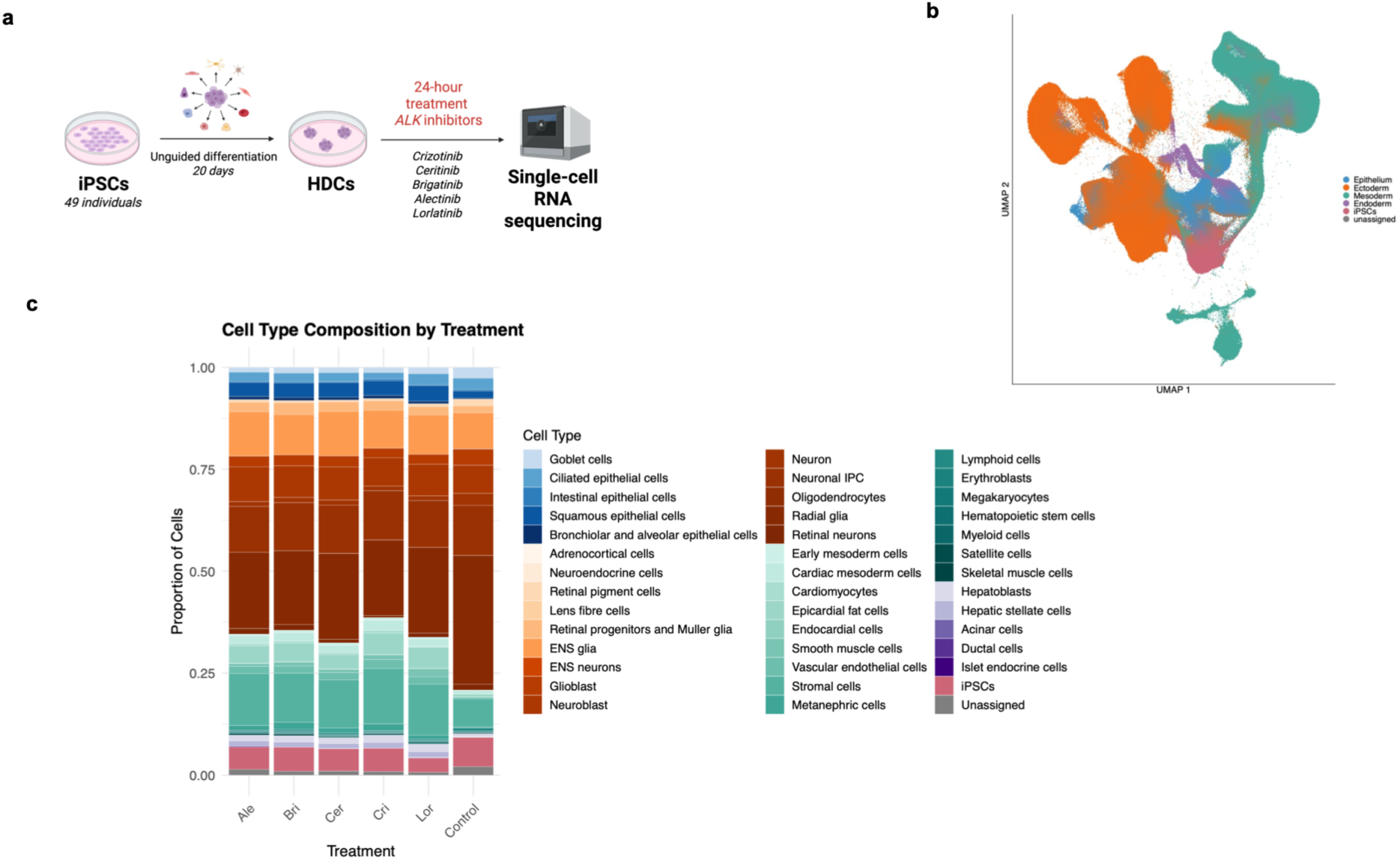
Single-cell RNA-sequencing of iPSC-derived heterogeneous differentiating cells treated with ALK inhibitors. (**a**) Experimental design: iPSCs from 49 donors were differentiated for 20 days to generate HDCs. HDCs were treated for 24 hours with each of five ALK inhibitors (alectinib, brigatinib, ceritinib, crizotinib, and lorlatinib), followed by scRNA-seq. (**b**) Uniform manifold approximation projection (UMAP) embedding of treated cells passing quality control across all 49 individuals and five conditions, colored by germ layer. (**c**) Cell type composition across treatment conditions. Stacked bars show the proportion of cells assigned to each of 41 cell types (Ale = alectinib; Bri = brigatinib; Cer = ceritinib; Cri = crizotinib; Lor = lorlatinib). Colors are grouped by germ layer family where blues = epithelium, oranges = ectoderm, teals = mesoderm, purples = endoderm, pink = iPSCs, and gray = unassigned.

After 24 hours of ALK inhibitor treatment, we dissociated the cells and prepared them for scRNA-seq on the 10X Genomics platform (Supplementary Data 1, Methods). After quality control, we retained data from a total of 1,962,313 cells with a median of 4,809 cells per sample and 4,595 unique molecular identifiers (UMIs) per cell (Supplementary Data 2, Methods).

Following the approach of Lin *et al.*, we used a cluster-free method to annotate cell types^34,35^ (Methods). While HDC cell types span all three germ layers, we observed ectodermal cell types to be the most abundant, reflecting the well-characterized tendency of iPSCs to differentiate into neuronal cells^36–38^ (Fig. 1b). Across the five treatment conditions, we annotated 41 cell types with a median of 14,068 cells per type (Supplementary Data 3). Consistent with previous studies using HDCs^32,34^, we observed between 31 to 41 cell types per individual (median of 38) across the five treatment groups (Supplementary Data 4; Supplementary Fig. 1). All 41 cell types were present in the control dataset, where we identified between 24 to 39 cell types per individual (median of 35; Supplementary Data 4).

The relatively consistent cellular composition of HDCs across conditions (Fig. 1c) suggests that the treatments did not affect differentiation in a way that substantially altered cell type composition or proportions. Thus, the transcriptional differences we observed between treatments in our downstream analyses may be reasonably interpreted as driven by actual transcriptional differences rather than differences in cell type composition.

### ALK inhibitors induce distinct transcriptional responses within and between cell types

Although all five drugs in this study are designed to inhibit ALK activity, they differ in their off-target profiles and tissue-specific side effects. To identify transcriptional effects that distinguish individual ALK inhibitors, we adopted a direct comparison across drugs rather than comparing each drug with untreated controls and then evaluating differences between their responses. Specifically, for each inhibitor, we compared gene expression in cells treated with that drug to the average expression across the other four drug-treatment groups^39^ (Methods).

Considering data from each cell type individually, we identified thousands of differentially expressed genes (DEGs) between each treatment and the average response to the other four inhibitors (Fig. 2a, Supplementary Data 5). Lorlatinib treatment resulted in the largest number of DEGs (66,155 DEGs across 30 cell types; 14,246 unique genes; FDR < 0.05), followed by alectinib (24,948 DEGs; 8,950 unique), crizotinib (15,091 DEGs; 6,277 unique), brigatinib (7,507 DEGs; 3,464 unique), and ceritinib (5,639 DEGs; 2,656 unique). The number of DEGs is correlated with the treatment concentration (Methods; 10 μM for lorlatinib, 1 μM for alectinib, 0.3 μM for crizotinib, 0.2 μM for brigatinib, 1 μM for ceritinib). We therefore do not draw conclusions based on the relative numbers of expression changes following the treatments. The proportion of DEGs discovered in each cell-type— treatment combination is also correlated with cell type abundance (Spearman ρ=0.82-0.95; p ≤2.4 x 10^−7^; Supplementary Fig. 2), and reflect, at least in part, differences in power.

**Figure 2.**
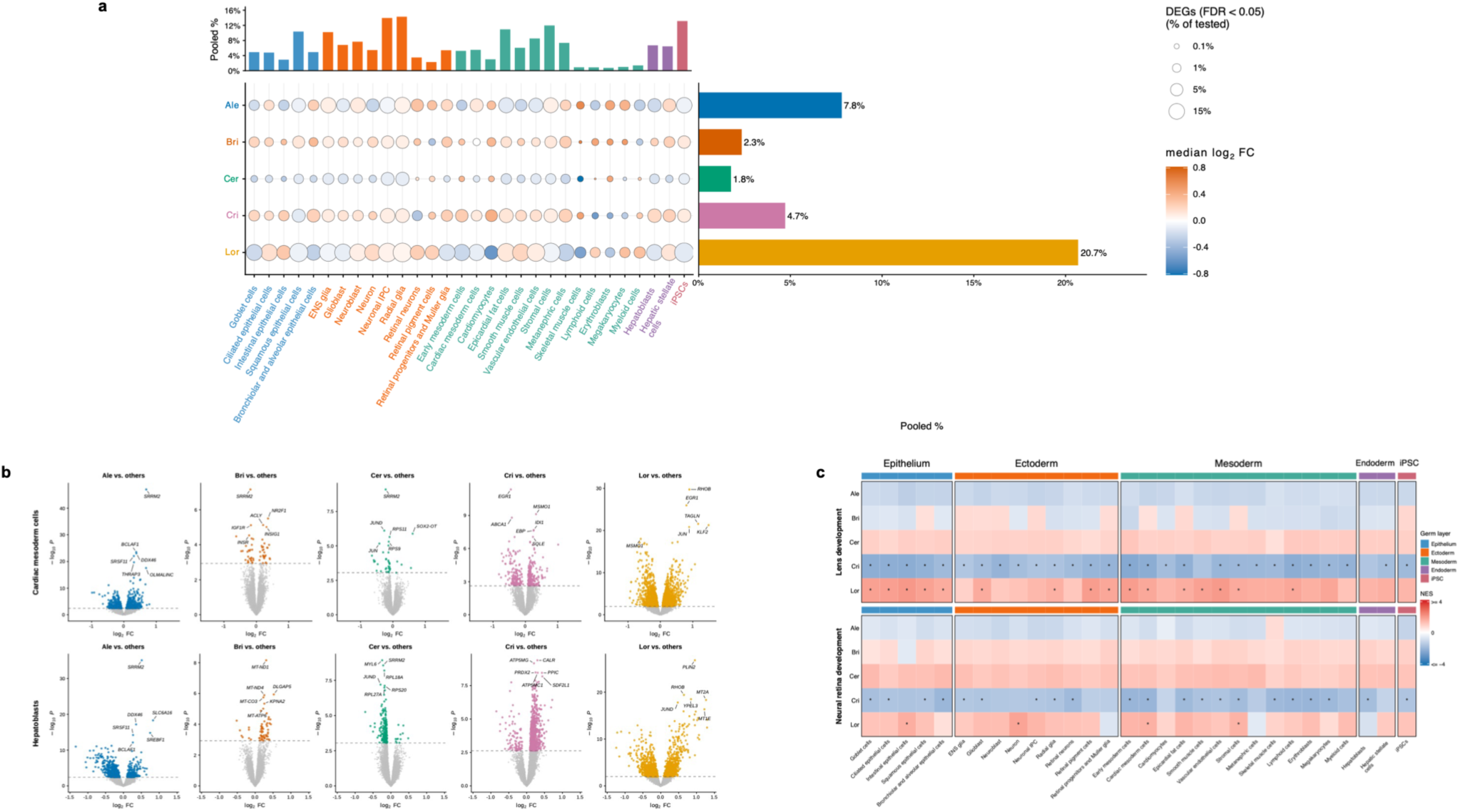
Shared and context-specific transcriptional responses to ALK inhibitors. (**a**) Dot plot of DEGs (FDR < 0.05) identified in each cell-type–treatment condition. Dot size denotes the percentage of significant DEGs out of all genes tested in the focal condition; dot color denotes the median effect size (log2 fold-change) among significant genes. Marginal bar plots show the pooled DEG fraction, or number of significant gene–cell-type–drug tests divided by the total number of tests performed across the five drugs for each cell type (top) or across the 30 cell types for each drug (right). Cell types are grouped by germ layer: epithelium (blue), ectoderm (orange), mesoderm (teal), endoderm (purple), and iPSCs (pink). (**b**) Volcano plots of DEGs in cardiac mesoderm cells (top) and hepatoblasts (bottom) treated with each of the five drugs. Significant DEGs (FDR < 0.05) are shown in color and the six genes with the lowest p-values are labeled. (**c**) Heatmap showing percell-type NES for two GO BP eye-development pathways (lens development and neural retina development) across celltype—treatment conditions, grouped by germ layer. * indicates p-adj < 0.05. *Abbreviations*: Ale, alectinib; Bri, brigatinib; Cer, ceritinib; Cri, crizotinib; Lor, lorlatinib; FC, fold-change; FDR, false discovery rate.

While many DEGs are shared by multiple cell types within the same treatment group (Supplementary Data 6), the overall transcriptional response to a given drug is distinguishable from one cell type to the next (Fig. 2b; results from other cell types are provided in Supplementary Fig. 3). To assess the extent of shared and specific DEGs among cell types treated with the same drug, we applied multivariate adaptive shrinkage (mash)^40^ to DEGs from each treatment group (Methods). By jointly estimating DEG effects across cell types, mash improves power to detect shared DEGs^40^. Among cell types treated with the same drug, we define a DEG as shared if its local false sign rate (lfsr) is less than 0.05 in at least two cell types. Conversely, we classified DEGs as cell-type-specific if they were tested in at least two cell types from a given treatment group, and if the gene’s lfsr is less than 0.05 in one cell type and either meets or exceeds 0.05 in all other tested cell types. This approach is conservative with respect to classifying sharing, and thus, as expected, the majority of DEGs are classified as shared; however, we identified 1,687 cell-type–specific DEGs associated with lorlatinib treatment, 1,605 with alectinib treatment, 1,314 with crizotinib treatment, 624 with brigatinib treatment, and 354 with ceritinib treatment (Supplementary Data 7, Supplementary Data 8; Supplementary Fig. 4).

To learn which biological pathways and processes are affected by each of the ALK inhibitors, we ranked genes in each cell-type—treatment condition using the statistics output by mash. We then applied fGSEA^41^ to perform gene set enrichment analyses (GSEA) using the MSigDB Hallmark pathways and GO Biological Process (GO BP) gene sets (Methods; Supplementary Figs. 5-7; Supplementary Data 9-10). To summarize the general, pathway-level effects of each ALK inhibitor, we calculated an average normalized enrichment score (NES) for each pathway tested under each treatment group (i.e., the mean NES across all tested cell types treated with the same drug). In the following section, we highlight observed patterns of enrichment and differential expression that are consistent with a drug’s known side effects or with its previously reported molecular phenotypes.

### Drug-induced transcriptional changes implicate candidate genes relevant to adverse events

When we considered individual DEGs and pathways affected by treatments with ALK inhibitors, we found several observations that may be related to reported toxicities. A full list is available in Supplementary Figs. 5-7 and Supplementary Data 9. Here we mention a few compelling examples.

Among the inhibitors we used in this study, crizotinib treatment is most frequently associated with visual disturbances (Table 1). In our data, DEGs with decreased expression in crizotinib-treated cells are enriched among genes involved in lens development (p-adj <0.05 in 28 out 30 cell types tested; mean NES = −2.06) and neural retina development (p-adj <0.05 in 21 out of 30 cell types tested, mean NES=−1.84) (Fig. 2c). We did not observe enrichment of these pathways when we considered DEGs with decreased expression in response to any other treatment. Moreover, in retinal cell types, we identified multiple cell-type-specific DEGs for crizotinib that have been linked to ocular abnormalities. In retinal progenitors/Müller glial cells we identified *ABHD12* (β=0.06, lfsr=0.037), which has been linked to retinitis pigmentosa and cataracts^42,43^, as well as *UBR7* (β=0.12, lfsr=0.015), whose ablation produces abnormal retinal morphology in mice^44^. Among DEGs specific to retinal neurons, *ELOVL2* (β=−0.16, lfsr=0.012) is associated with retinitis pigmentosa and macular degeneration in mice^45,46^, *SDCCAG8* (β=−0.22, lfsr=0.008) has been linked to retinal-renal ciliopathy^47^, and *GTDC1* (β=−0.19, lfsr=0.037) is associated with corneal astigmatism^48^ and myopia^49^.

Hepatotoxicity is the most commonly-reported adverse event in patients treated with alectinib (Table 1). We identified 67 cell-type-specific DEGs in alectinib-treated hepatoblasts, including *ATOX1* (β=−0.24, lfsr =5.8×10^−4^), which was not a DEG in any of the other 29 alectinib-treated cell types (lfsr ≥ 0.97). *ATOX1* (also known as *HAH1*) encodes a cytosolic copper chaperone and antioxidant that delivers copper to ATP7B, an ATPase required for biliary copper excretion in hepatocytes^50^. Other hepatoblast-specific DEGs in the alectinib group include the copper- and zinc-binding metallothionein genes *MT1E, MT1F, MT1M,* and *MT1X*, which are important for regulating oxidative stress^51^. Like *ATOX1*, all four genes are downregulated in response to alectinib (mean β=−0.56; lfsr<0.05; Supplementary Data 8), suggesting that alectinib-induced hepatotoxicity could be mediated by a reduced capacity to maintain copper homeostasis or regulate antioxidant activity.^52,53^

Ceritinib treatment is associated with hyperglycemia (Table 1), which has been suggested to reflect reduced insulin sensitivity^54–56^. Ceritinib is also the only ALK inhibitor in this study whose FDA label identifies the insulin receptor (which mediates the metabolic effects of insulin, including glucose uptake) as a target at clinically relevant concentrations^57^. We found that DEGs with decreased expression in ceritinib-treated cells are enriched among genes involved in cytoplasmic translation (p-adj<0.05 in all 30 tested cell types, mean NES =−3.05) and ribosome biogenesis (p-adj<0.05 in all 30 tested cell types; mean NES = −2.62). These genes include *RPS9* and *RPL27A*, which encode ribosomal proteins and had decreased expression in ceritinib-treated cells of all 30 cell types (mean β =−0.11 and −0.12, respectively; lfsr < 1.0 x 10^−7^ in all cell types). Insulin signaling promotes both cytoplasmic translation and ribosome biogenesis^58^, in part through activation of mTORC1^59,60^. Consistent with reduced insulin signaling, the mTORC1 signaling Hallmark pathway was downregulated in response to ceritinib treatment (mean NES = −1.70, p-adj. < 0.03 in all cell types).

Lorlatinib treatment is associated with hyperlipidemia^61^ (Table 1) and disruptions in lipid metabolism and secretion^61^. In clinical trials, lorlatinib treatment was specifically associated with increase in triglyceride and total cholesterol levels^26^. Consistent with these reports, we found that DEGs showing increased expression in lorlatinib-treated cells are enriched for lipid-related gene sets, including regulation of plasma lipoprotein particle levels (p-adj <0.05 for 26 out of 29 tested cell types; mean NES = 2.06). We also found the cholesterol homeostasis pathway to be enriched among DEGs showing decreased expression in lorlatinib-treated cells (p-adj <0.05 in 29 out of 30 cell types tested; mean NES = −2.37; Fig. 2c), with sterol biosynthetic process genes following a similar pattern (p-adj < 0.05 for 28 out of 30 tested cell types, mean NES = −2.14). The decreased expression of sterol synthesis genes, which are suppressed by sterol accumulation^62^, are corroborated by previous reports that lorlatinib promotes intracellular cholesteryl-ester accumulation^63^.

A gene-specific observation that may be relevant in the context of liver injury relates to the effect of lorlatinib treatment on the expression of *PLIN2*, particularly in hepatoblasts (β = 0.87, lfsr = 4.86×10^−40^; see Fig. 2b), epicardial fat cells (β = 0.79, lfsr = 3.07×10^−75^), and hepatic stellate cells (β = 0.69, lfsr = 8.06×10^−23^). *PLIN2* encodes the lipid droplet-binding protein Perilipin-2. Elevated expression of Perilipin-2 results in increased fatty acid uptake; moreover, *PLIN2* has been associated with a variety of metabolic traits, including fatty liver disease, heart disease, and type 2 diabetes^64^.

Among the five ALK inhibitors included in this study, lorlatinib was specifically designed for improved penetration of the blood-brain barrier. Accordingly, it has distinct effects on the CNS, producing a range of neurocognitive and mood effects^25^ in approximately 52% of patients (Table 1). In our data, lorlatinib treatment is exclusively associated with increased expression of *GRIA2* in ectodermal-derived cell types (neuronal IPCs β=0.58, lfsr=2.02×10^−10^; radial glial cells β=0.40, lfsr=5.83×10-^6^; ENS glial cells β=0.33, lfsr=1.37×10^−4^; neurons β=0.27, lfsr=1.23×10^−2^; and retinal neurons β=0.26, lfsr=2.11×10^−2^). These observations, across multiple related cell types, suggest that lorlatinib treatment may have distinct lineage-dependent effects on *GRIA2* expression (Supplementary Data 9). *GRIA2* encodes an AMPA receptor subunit crucial for post-synaptic signaling and long-term potentiation^65^, and mutations in GRIA proteins have been implicated in a variety of neurological abnormalities (e.g., epilepsy, intellectual disability, and sleep disorders)^66^.

Lorlatinib treatment is also associated in our data with increased expression of multiple genes encoding neurofilament proteins (*NEFM, INA*, *NEFH*, *NEFL*, and *PRPH*) compared to the other ALK inhibitors. These proteins facilitate intra-cellular transport and provide structural support for neurons^67^ (Supplementary Data 9). Treatment with other inhibitors either did not affect the expression of these genes or resulted in decreased expression levels (Supplementary Data 9). The neurofilaments encoded by these genes form heteropolymers that co-assemble in various combinations depending on the cell type and developmental stage^67^, which could explain some of the heterogeneity we observed among lorlatinib-treated cells. Because neurofilaments are released into cerebrospinal fluid and plasma as a result of neuronal damage, they are used as biomarkers for monitoring patients with traumatic brain injury, stroke, dementia, and other neurological disorders^68^.

### Mapping response eQTLs to treatments with specific ALK inhibitors

Genetic factors contributing to inter-individual variability in the transcriptional response to ALK inhibitors may be useful for predicting a patient’s risk for drug-induced adverse events, particularly if they regulate gene expression in tissues relevant to a drug’s commonly reported side effects (Table 1). We were therefore interested in identifying response eQTLs, where genetic effects on gene expression change with treatment, reflecting direct or indirect interactions between genotype and drug exposure.

As a first step, we mapped *cis* eQTLs separately in each cell-type–treatment condition, including untreated controls. We restricted our analysis to the 21 cell types represented by at least 20 individuals in every condition (excluded cell types are listed in Methods). After filtering and normalizing the data, we used linear models implemented in MatrixEQTL^69^ to test associations between gene expression and common SNPs (MAF > 0.05) within 100 kb upstream or downstream of each gene’s transcription start site (TSS). We included sex and gene expression principal components as covariates (Methods).

We then applied mash jointly across all treatment and control conditions to improve power to find eQTL by sharing information across contexts^40^. Using this approach, we classified 4,371 eGenes, namely genes associated with at least one cis-eQTL, in at least one context, at local false sign rate (lfsr) below 0.05 (Supplementary Data 11). Associations with 221 of these eGenes (5.1%) met this threshold in every condition, whereas associations with 990 eGenes (22.7%) met the threshold in exactly one condition. We detected a median of 1,328 eGenes per condition (Supplementary Fig. 8). The number of eGenes detected was positively correlated with the number of individuals represented in each condition (Pearson r = 0.69) and the number of cells (Pearson r = 0.78; Supplementary Fig. 9), consistent with differences in statistical power across conditions.

Our primary interest, as mentioned, was specifically to identify response eQTLs, namely loci whose effects on gene expression change in response to treatment. Differences in whether an association meets a significance threshold are insufficient to identify such loci: incomplete and uneven statistical power can make a shared effect appear treatment-specific or obscure a true difference between conditions. A straightforward approach to find response eQTLs is to explicitly test genotype-by-treatment interactions (Methods). We identified 173 loci with evidence for an interaction after multiple-testing correction (adjusted p < 0.05; Methods). However, formal interaction tests have limited power, especially given our sample size. We therefore also used an ad hoc approach applied in our previous work^34^, relying on mash effect-size estimates to identify marked differences in genetic effects on gene expression between treatment and control conditions.

For each cell type, we compared the mash posterior mean effect of each SNP–gene pair between each drug-treatment group and the untreated control group. We classified an association as a response eQTL if the lfsr was below 0.05 in at least one of the two conditions and the estimated effect sizes differed either in direction or by at least three-fold in magnitude. This definition combines statistical evidence for a nonzero genetic effect in at least one condition with a substantial difference in the estimated effect between conditions. Applying the effect-size criterion to mash posterior means reduces reliance on noisy estimates from separate per-condition regressions.

For loci supported by the interaction tests, the interaction coefficients were correlated with the corresponding differences in mash posterior mean effects between treatment and control conditions (Pearson r = 0.52–0.56 across the five drugs), with agreement in direction for 74.2–75.9% of loci (Supplementary Fig. 10). Using the mash-based approach, however, we identified response eQTLs for 2,137 genes, representing 48.9% of the 4,371 eGenes (Supplementary Data 12; Fig. 3a).

**Figure 3.**
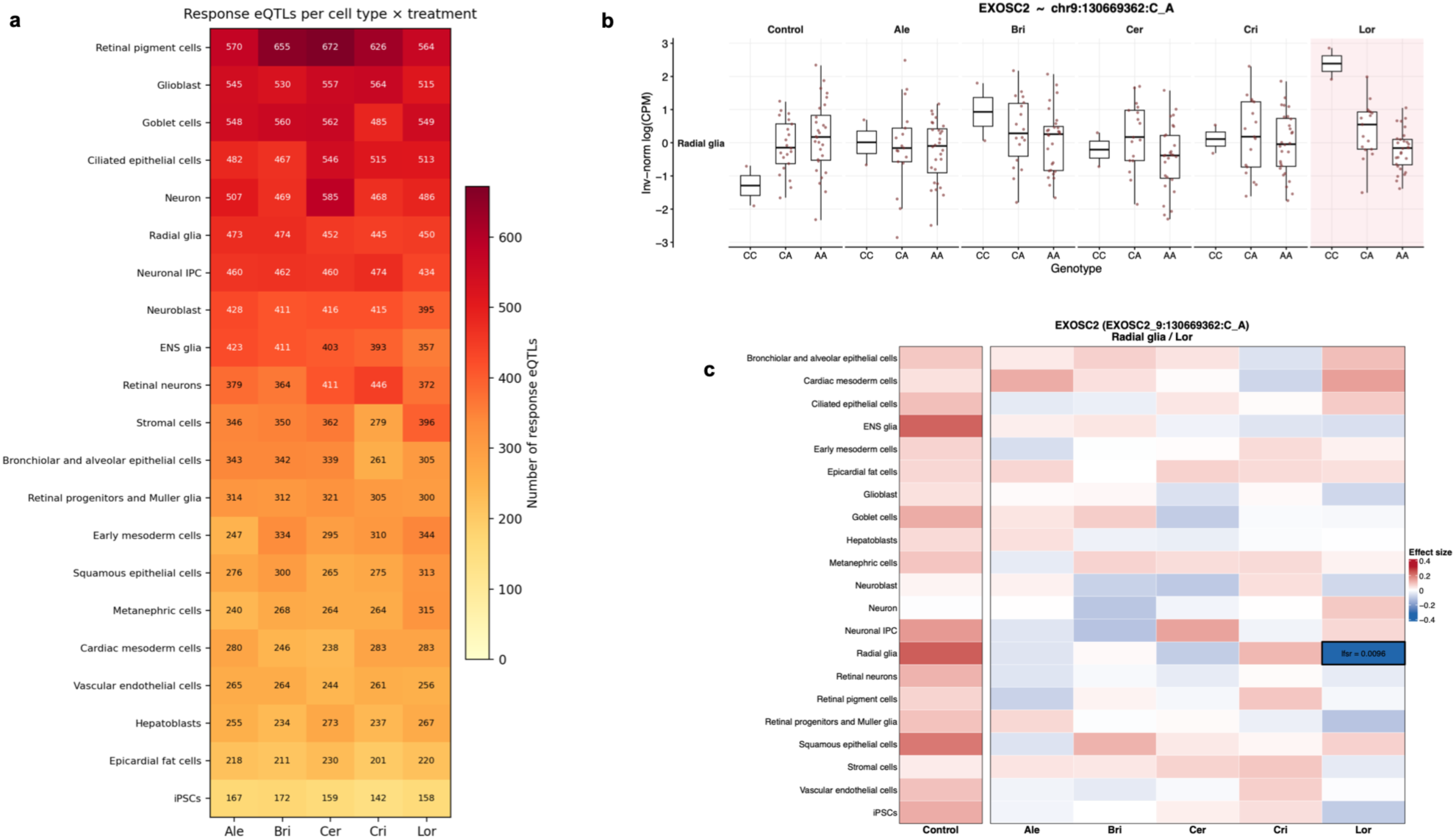
Response eQTL mapping identifies treatment-specific and cell-type-specific *cis*-regulatory effects. **(a)** Number of unique response eGenes discovered per cell type (rows) and treatment (columns). The same gene may be a response eGene in multiple conditions. **(b)** A response eQTL (chr9:130669362:C>A) specific to lorlatinib-treated radial glial cells is highlighted in pink. Box plots show the inverse-normal-transformed log(CPM) expression of *EXOSC2* by genotype in control and drug-treated radial glial cells, delimited by the inter-quartile range and split at the median. Each point is the expression level of *EXOSC2* averaged across all cells from a single individual in that condition and genotype class. **(c)** Mash posterior mean effect sizes (β) for the same eQTL across all conditions. Color indicates the direction and magnitude of the effect (red = positive, blue = negative). Lorlatinib-treated radial glial cells are distinguished by a black-outlined box.

Compared with non-response eQTLs, response eQTLs were less concentrated around TSSs, more enriched in proximal and distal enhancer-like signatures and depleted in promoter-like signatures (Supplementary Fig. 11; Supplementary Fig. 12; Supplementary Data 13). Response eGenes also had longer cumulative enhancer lengths and tended to be more intolerant to loss-of-function mutations (Methods; Table 3), consistent with previous reports^70^. These patterns were robust to increasing the effect-size threshold used to classify response eQTLs (Supplementary Data 14).

**Table 3.** Regulatory features of response eGenes vs non-response eGenes across treatment–cell-type conditions. Each row corresponds to one gene-level feature. Within each condition, we computed the feature value (proportion or mean) separately for response and non-response eGene sets and took the difference. The sign test counts the number of conditions in which response > non-response and tests against a null of 0.5 (one-sided binomial). FDR was BH-corrected across computed features.

| **Feature** | **Conditions with response > non-response**<br>N / total (%) | **Median change (response – non-response)** | **Sign-test p (one-sided)** | **BH FDR** |
|---|---|---|---|---|
| Proportion LoF-intolerant (pLI >=0.9) | 65/105 (62%) | 0.029 | 0.0094 | 0.047 |
| Mean TSS count per eGene | 60/105 (57%) | 0.246 | 0.086 | 0.086 |
| Mean cumulative ABC enhancer length per eGene | 63/105 (60%) | 20.3 | 0.025 | 0.038 |

The response eQTLs identified here capture interactions between genotype and drug treatment and provide a resource for identifying genetic predictors of individual susceptibility to ALK inhibitor toxicity. Establishing their predictive utility will require studies in patient cohorts. Below, we examine a subset of these response eQTLs to explore their potential mechanistic relevance to the adverse effects associated with individual ALK inhibitors.

### Response eQTLs are present in cell types representing tissues affected by ALK inhibitor toxicities

We found multiple response eQTLs with plausible connections to the adverse events associated with ALK inhibitors (we provide a complete list of annotated response eQTLs in Supplementary Data 15). Given that ALK inhibitors are associated with a variety of adverse events, we leveraged information from the drugs’ FDA labels to prioritize response eQTLs from cell types that could plausibly be connected to adverse events affecting specific tissues (Table 1, Supplementary Data 15). The examples discussed below are summarized in Table 4; we also provide a comprehensive list of response eQTLs per treatment, cell type, and condition in Supplementary Data 12.

**Table 4.** Examples of response eQTLs in cell types relevant to known ALK inhibitor toxicities.

| **Drug** | **Adverse event** | **Cell type** | **Variant** | **eGene** | **Control β (lfsr)** | **Treatment β (lfsr)** | **Function** |
|---|---|---|---|---|---|---|---|
| Crizotinib | Visual disturbances | Retinal neurons | chr17:7234631C>G | *ELP5* | 0.81 (6.0×10<sup>-8</sup>) | 0.25 (0.035) | Elongator complex subunit; retinal ganglion cell survival |
|  | Visual disturbances | Retinal neurons | chr1:236067600G>A | *NID1* | –0.06 (0.29) | –0.23 (0.0044) | Basement-membrane glycoprotein; anti-angiogenic retinal role |
| Alectinib | Hepatotoxicity | Hepatoblasts | chr8:11916946A>G | *CTSB* | –0.034 (0.344) | -0.127 (0.012) | Lysosomal protease; hepatocyte apoptosis |
| Brigatinib | Photosensitivity/rash | Squamous epithelial cells | chr2:38053953C>G | *CYP1B1* | 0.05 (0.31) | 0.21 (0.007) | UV-induced cytochrome P450; drug phototoxicity |
|  | Visual disturbances | Retinal pigment cells | chr15:43187130C>T | *CCNDBP1* | 0.436 (3.0×10<sup>-6</sup>) | 0.140 (0.031) | Cyclin D1/CDK4 regulator; retinal progenitor proliferation |
|  | Hypertension | Vascular endothelial cells | chr5:80375422C>A | *ZFYVE16* | 0.001 (0.493) | 0.133 (0.036) | BMP/SMAD signaling adaptor; endothelial pathobiology |
| Ceritinib | GI toxicity | Goblet cells | chr9:39267725C>T | *CNTNAP3* | –0.028 (0.320) | –0.125 (0.033) | Linked to intestinal epithelial autophagy (ATG16L1) |
| Lorlatinib | CNS effects | Radial glia | chr9:130669362C>A | *EXOSC2* | 0.239 (0.082) | –0.318 (0.0096) | RNA exosome cap subunit; neurological disease |
|  | CNS effects | Neurons | chr9:137286910C>G | *NRARP* | 0.025 (0.411) | –0.118 (0.039) | Notch pathway feedback regulator; neural progenitor maintenance |

In radial glial cells, we identified a lorlatinib response eQTL for *EXOSC2,* which encodes a component of the RNA exosome. *EXOSC2* is also a DEG with decreased expression levels in six lorlatinib-treated cell types, including radial glial cells (β =−0.049, lfsr =0.019). In untreated radial glia this locus is not identified as an eQTL, with the effect on expression revealed only after treatment with lorlatinib, but not with any of the other drugs (β = −0.32, lfsr = 9.6 x 10^−3^; Fig. 3b-c). In addition to their role as neural progenitors, radial glial cells regulate synaptic plasticity in the adult cerebellum. *EXOSC2* polymorphisms have been linked to neurological diseases, including rare syndromes marked by cerebellar degeneration and atrophy^71,72^. This response eQTL therefore is a candidate biomarker for inter-individual risk for the known adverse CNS effects associated with lorlatinib.

Crizotinib is commonly associated with visual disturbances, with all-grade reported frequencies of 60-71%^73^ (Table 1). In vision-relevant cell types (retinal pigment cells, retinal neurons, and/or retinal progenitors and Müller glia), across all treatments, we identified 83 response eQTLs (Supplementary Data 16). One of these response eQTLs is specific to crizotinib-treated retinal neurons, where it is associated with the expression of *ELP5,* a gene encoding an Elongator subunit involved in tRNA binding^74,75^. This eQTL is detected in both untreated and treated cells, but the effect size is markedly different following the treatment (β = 0.81 before, β = 0.25 after). Retinal loss of a related Elongator subunit, ELP1, has been shown to cause mitochondrial dysfunction and selective retinal ganglion cell degeneration; moreover, retinal ganglion cell loss has been observed in familial dysautonomia patients with Elongator pathway mutations^76,77^. Elongator-dependent mitochondrial vulnerability may therefore be modified by genotype-dependent regulation of *ELP5* in crizotinib-treated retinal neurons.

We also found a crizotinib response eQTL in retinal neurons for *NID1*, which encodes Nidogen-1, a glycoprotein supporting the ocular basement membranes and retinal vasculature^78^. Nidogen-1 has previously been linked to diabetic retinopathy^79^ and subretinal neovascularization (abnormal growth of blood vessels in the choroid layer of the eye) which can cause visual disturbances and in many cases, precedes age-related macular degeneration^78^. In a mouse model of subretinal neovascularization, *Nid1* knockout mice show larger, leakier choroidal neovascular lesions compared to control mice, consistent with a role for NID1 in stabilizing an anti-angiogenic retinal basement membrane^78^.

While all five ALK inhibitors are associated with symptoms of liver dysfunction, hepatotoxicity is particularly common with alectinib treatment, occurring in 41% of patients, with severe (Grade 3-4) adverse events in 8% of treated patients (Table 1). In hepatoblasts, we identified an alectinib response eQTL for *CTSB*. The eQTL is not detected in untreated cells or cells treated with the other drugs. *CTSB* encodes Cathepsin B, a lysosomal cysteine protease with a role in hepatocyte apoptosis^80^. It was previously linked to hepatotoxicity induced by vandetanib, a kinase inhibitor that, like alectinib, has activity against RET (Table 1); specifically, drug-induced STAT3 activation was shown to upregulate transcription of *CTSB*, triggering hepatocyte apoptosis^81^. Further, *Ctsb* inactivation reduces hepatocyte apoptosis and liver injury across several mouse models of cholestatic and ischemic liver damage^82,83^. Together, these observations suggest that genotype-dependent regulation of *CTSB* under alectinib treatment could influence susceptibility to hepatotoxicity.

Gastrointestinal toxicity occurs frequently with ceritinib. Diarrhea, nausea, vomiting, or abdominal pain occurred in 95% of patients across clinical studies, with 14% of patients experiencing severe (Grade 3-4) adverse events (Table 1). We identified a response eQTL for *CNTNAP3* that is specific to ceritinib in goblet cells, which produce mucus^84^. *CNTNAP3* encodes a member of the contactin-associated protein family of neurexin-related cell-recognition molecules^85^; this gene is reportedly upregulated in intestinal tissue from Crohn’s disease patients, where it increases expression of the autophagy gene *ATG16L1* (a Crohn’s disease risk gene) and promotes autophagosome formation^86^. Autophagy has been connected to gastrointestinal disease, including inflammatory bowel disease and functional dyspepsia, through its role in goblet cell homeostasis, balancing mucus production and ER stress^87,88^. A ceritinib-specific change in *CNTNAP3* expression in goblet cells could potentially be relevant for altered intestinal epithelial autophagy related to gastrointestinal symptoms.

While dermatological side effects (i.e., photosensitivity and rash) have been reported following treatment with a number of ALK inhibitors, they are especially serious for brigatinib. Approximately 0.7% of patients treated with brigatinib experience severe (Grade 3-4) adverse events, which is sufficient to justify a modified treatment plan (Table 1). We identified a brigatinib-specific eQTL for *CYP1B1* in squamous epithelial cells. We did not identify this locus as a response eQTL in any other cell-type—treatment condition. *CYP1B1* encodes the most abundant cytochrome P450 in human skin and is induced in keratinocytes by UV-B ^89,90^. *CYP1B1* genotypes have been shown to modulate cutaneous sensitivity to drug phototoxicity in the psoralen-UVA system^91^, making a genotype-dependent, drug-induced change in cutaneous CYP1B1 a plausible marker for individual variability in photosensitivity.

Brigatinib is also associated with visual disturbances, with 7.4% of patients experiencing Grade 1-2 adverse events (Table 1). We identified a brigatinib response eQTL only in retinal pigment cells for *CCNDBP1* (cyclin D1 binding protein 1) (Supplementary Data 17). *CCNDBP1* negatively regulates the cyclin D1/CDK4 complex and is associated with optic disk drusen and branch retinal artery occlusion in genetic disease databases^92^. Cyclin D1 is the predominant cyclin in the developing retina; mice that lack it have small eyes and hypocellular retinas due to reduced retinal progenitor proliferation^93,94^.

Finally, we identified a brigatinib-specific response eQTL for *ZFYVE16* in vascular endothelial cells, a cell type that is plausibly connected to brigatinib-associated hypertension (32% all-grade; Table 1). *ZFYVE16* encodes Endofin, an endosomal FYVE-domain adaptor that promotes phosphorylation of SMAD1 and acts as a SMAD anchor for BMP receptor signaling and binding^95^. *ZFYVE16* has been nominated as a risk gene for sporadic brain arteriovenous malformation, in which BMP/TGF-β signaling is also implicated^96^. Given the established contribution of endothelial BMP-SMAD1 signaling to the pathobiology of pulmonary arterial hypertension^97^, brigatinib’s associated vascular side effects could be mediated in part by gene regulatory changes to this endothelial BMP-SMAD adaptor in vascular tissue.

## Discussion

ALK-positive non-small cell lung cancer (NSCLC) presents a challenge for selecting treatments based on both efficacy and tolerability. Several ALK inhibitors provide effective therapeutic options, but their adverse-event profiles differ substantially. For patients with more than one clinically appropriate option, differences in toxicity may be more relevant to treatment choice than differences in expected efficacy. Severe adverse events may lead to dose modifications, treatment interruptions, or a switch to another inhibitor after treatment has begun. Identifying patients at increased risk for specific toxicities before treatment could reduce the need for such changes.

Although the adverse events associated with individual ALK inhibitors are well documented, the transcriptional responses of non-cancer cells to these drugs have not been systematically characterized. Consequently, little is known about which genes, pathways, and cell types may contribute to their different toxicity profiles. It is also unclear why susceptibility to a particular adverse event differs among patients treated with the same drug. Genetic variation could contribute to this heterogeneity, but its role remains largely unexplored. The relatively small number of patients with ALK-positive NSCLC, combined with the diversity of drug-specific adverse events, makes well-powered genetic association studies difficult. In particular, there are no established GWAS loci for ALK inhibitor-associated toxicities to guide systematic investigation of their genetic basis.

In this study, we took first steps towards identifying genes, pathways, and genotypes that may be relevant to individual responses to ALK inhibitors. By treating heterogeneous differentiating cultures (HDCs) derived from 49 genotyped individuals with five inhibitors, we characterized transcriptional responses across diverse non-cancer cell types. We found that drugs sharing the same primary target nevertheless produced distinct transcriptional responses, often in a cell-type-specific manner. Our one-versus-four analysis choice allowed us to identify effects specific to each drug, which a treatment-versus-control analysis would not be able to distinguish from a class-shared ALK effect. This is particularly relevant because differences between ALK inhibitors may contribute to their distinct side effect profiles. Some of the affected genes and pathways in retinal, hepatic, neural, intestinal, and vascular cell types are consistent with the tissues in which individual inhibitors cause adverse events. These observations do not establish mechanisms of toxicity; instead, they allow us to formulate biological hypotheses that can be pursued in future studies of the corresponding adverse events.

The genetic analysis adds a second dimension to this resource. We identified response eQTLs involving 2,137 genes across 21 cell types. These associations identify genotypes whose effects on gene expression differ between drug-treated and untreated cells, potentially reflecting direct or indirect interactions between genotype and drug exposure. An association between genotype and a transcriptional response, however, is not evidence that the same genotype predicts toxicity in patients. Each response eQTL therefore nominates a locus at which genetic variation may modify the transcriptional response to a particular ALK inhibitor in a defined cell type. Together with the drug-specific transcriptional profiles, these associations can help prioritize candidate mechanisms and genetic markers for particular adverse events (Table 4).

The central limitation of this study follows directly from the gap that motivated it. Without GWAS loci associated with ALK inhibitor toxicities, we could not test whether the response eQTLs share genetic association signals with clinical outcomes through colocalization. Existing GWAS for broader traits, such as hypertension, do not necessarily provide adequate substitutes, because drug-induced adverse events may have different genetic determinants. At this time, we also lack patient data with which to test whether the variants identified in HDCs are associated with adverse events. Consequently, we cannot establish causal relationships between these regulatory associations and clinical toxicity, assign risk to individual genotypes, or evaluate their predictive performance.

The immediate deliverable from this study is a searchable resource for developing focused hypotheses about ALK inhibitor toxicity. Retrospective cohorts with genotypes and well-characterized adverse events, or prospective studies designed to collect such data, could test whether selected response eQTLs are associated with the specific toxicities of their corresponding drugs. A recent retrospective study of 215 patients treated with alectinib found that homozygous carriers of a common variant in *PPARA*, which encodes a transcription factor that regulates the drug-metabolizing enzyme CYP3A4^98^, had higher alectinib exposure and a higher incidence of grade 3 or higher toxicity.^99^ The variants tested in that study were selected in advance based on their known pharmacokinetic effects on drug-metabolizing enzymes and transporters. The response eQTLs we report here could eventually serve as additional candidates for studies of similar design. Functional perturbation of candidate regulatory variants or genes in appropriate cell models could provide complementary evidence about mechanism. We provide the cell-type-specific transcriptional results and response eQTLs for further investigation through an interactive viewer (https://wooeg.shinyapps.io/alk-resource/).

Eventually, these candidates biomarkers could be incorporated (in combination with clinical data) into classifiers to predict an individual patient’s likelihood of developing adverse events in response to a given drug. This could potentially inform treatment decisions. For example, if a patient has a high likelihood of developing a severe adverse event with one ALK inhibitor, a clinician may consider using an alternative ALK inhibitor. Any eventual clinical predictor would also need to be evaluated alongside antitumor efficacy, resistance mutations, and other factors that influence drug choice.

An agnostic search for genetic predictors or biomarkers of individual ALK inhibitor toxicities is unlikely to be well powered in the small patient cohorts available. The response eQTLs and drug- and cell-type-specific transcriptional responses identified here allow us to nominate candidate loci, genes, and pathways with plausible mechanistic links to specific adverse events. These biologically informed candidates can guide focused genotype-toxicity tests in retrospective and prospective patient cohorts, providing a more tractable route toward developing biomarkers of individual susceptibility.

## Methods

### Selection of ALK inhibitor doses and treatment duration

To select an appropriate treatment dose for each drug, we performed a pilot study. We consulted existing literature to select a range of three concentrations to test in three HDC lines. Because it is common for *in vitro* systems to require higher concentrations than would be expected based on the *in vivo* human C_max_, we tested concentrations that have been used in both *in vivo* and *in vitro* studies. On day 21 of HDC culture, we separately treated each line with each of the three concentrations selected for alectinib (MedChemExpress HY-13011A), brigatinib (MedChemExpress HY-12857), ceritinib (MedChemExpress HY-15656A), crizotinib (MedChemExpress HY-50878A), and lorlatinib (MedChemExpress HY-12215). At 24 and 48 hours, we used a microscope to assess cell viability, noting changes in cell morphology and the level of cellular debris. We selected the highest drug concentration at which all three HDCs were viable (minimal cellular debris and signs of cell death) after 24 hours to use in our eQTL study, which was 1 μM for alectinib, 0.2 μM for brigatinib, 1 μM for ceritinib, 0.3 μM for crizotinib, and 10 μM for lorlatinib. We chose a 24-hour treatment duration because we expected that a longer treatment duration would be more likely to impact cell type proportions within HDCs (e.g., by altering rates of cellular differentiation or causing rare cell types to drop out).

### HDC generation and treatment

Following the unguided differentiation protocol described by Popp *et al*., we generated HDCs from an existing panel of iPSCs derived from 30 female and 19 male Yoruba individuals from Ibadan, Nigeria (YRI)^33^. All individuals were fully genotyped using 30X whole-genome-sequencing data collected for the 1000 Genomes Project^100^. We differentiated iPSCs into HDCs in four independent batches, with each batch containing both male and female lines. The first three batches included 14-16 individuals each, and a fourth batch included 19 individuals, 14 of which had also been collected in an earlier batch to improve their representation (Supplementary Data 2). On day 20, within each batch, we pooled cells from all individuals in equal proportions in a single tube and then divided the cells into five tubes, one for each of the five ALK inhibitor treatments. We individually treated HDCs with 1 μM alectinib, 0.2 μM brigatinib, 1 μM ceritinib, 0.3 μM crizotinib, and 10 μM lorlatinib for 24 hours. After treatment, we dissociated the cells and prepared scRNA-seq libraries using the 10X Genomics 3’ HT Reagent Kit (v3).

### Single-cell RNA-sequencing

For each batch, we loaded each combined donor-treatment pool across two wells of a 10X Genomics chip, targeting ∼5,000 singlets per donor. In a previous eQTL mapping study, Popp *et al.* collected scRNA-seq data that included untreated HDCs derived from 46 out of 49 iPSC lines included in the current study^32^. Since our primary goal was to map *cis-*eQTLs, and *cis* genetic variation is unlikely to correlate with batch or other technical effects, we used these data as a control group for identifying drug response eQTLs, as we describe in detail below. We note that both the ALK inhibitor dataset and Popp *et al.* dataset were generated in the same laboratory by the same personnel, using the same differentiation protocol and iPSC panel, the 10X Genomics platform, and Illumina sequencing instruments.

scRNA-seq libraries were sequenced by the University of Chicago Functional Genomics Core on an Illumina NovaSeq X Plus instrument (28/88 bp paired-end) at a target depth of 15,000 reads per cell. This depth was chosen with the goal of discovering eQTLs with the strongest effects (i.e., those most likely to eventually be informative for a patient classifier) rather than maximizing total eQTL discovery. We aligned samples to the human genome (GRCh38) using 10X CellRanger v7.2.0^101^. We then used Vireo to demultiplex samples by genotype and assign droplets to individuals^102^ using a reference panel from 1000 Genomes Project Phase 3^100^. Using Seurat v5.0.2^103^, we removed doublets and droplets that could not be confidently assigned to an individual. We filtered cells to keep only those with percent mitochondrial reads less than 20% and with 1000-8000 unique genes detected in each cell. Following preliminary quality control, we retained 1,962,313 high-quality cells (median 4,595 UMIs and 2,459 genes detected per cell; median 2.54% mitochondrial reads) across the 49 donors.

For the control HDC dataset collected by Popp *et al.*^32^, we applied the preliminary quality control thresholds reported in their paper for their data. We filtered out doublets and unassigned cells, retaining cells with <15% mitochondrial reads and with at least 2,500 unique genes per cell. We removed cells with total UMI counts over 150,000 and required that genes be expressed in at least 10 cells. These thresholds differ from those applied to the treatment data primarily because of the difference in sequencing depth (Popp *et al.* targeted a sequencing depth of 100,000 reads per cell). We restricted the control data to the 46 YRI donors that were included in the treatment panel, resulting in 901,966 cells after quality control. All downstream eQTL analysis steps were performed identically in the control and treatment conditions.

### Cell type annotation

We annotated cell types using a previously validated method described by Lin *et al*.^31^. We used the CelliD annotation approach, which identifies gene signatures for each cell rather than relying on clustering^35^. Briefly, we used a combined reference list of cell type markers from the human fetal cell atlas^104^ and first-trimester fetal brain atlas^105^ to annotate the cells belonging to each batch–treatment combination; we then merged the annotated data for downstream analyses. Using this pipeline, we mapped 1,942,953 treated cells to 41 cell types with 19,360 cells remaining unassigned. We also annotated untreated HDCs from Popp *et al*. using the same pipeline, assigning 895,673 cells to 41 cell types with 19,103 cells unassigned (Supplementary Data 3).

To enable UMAP visualization of the full single-cell treatment dataset, we performed dimensionality reduction using the *SketchData* function, as described in Hie et al. 2019^106^. We used BPCells (v.0.1.0) to write the post-quality-control count matrix to an on-disk format^107^ and sub-sampled 850,000 cells from the full dataset using Seurat v5.0.2 Leverage Score sketching. We log-normalized counts in the sketched cells and used a variance-stabilizing transformation to identify 3,000 variable features. Following scaling, we performed principal component (PC) analysis on the sketched assay. We generated the UMAP using the first 30 PCs and projected the UMAP embedding onto the full dataset.

### Differential expression analysis

To identify transcriptional responses that differ between *ALK* inhibitors, we performed differential expression analysis using dreamlet (v.1.9.2)^39^. Since we did not include a within-experiment vehicle control, we used a one-versus-four (1v4) treatment contrast in which we compared, for each cell type, the expression of a gene in one treatment group to the gene’s average expression across the other four treatment groups (e.g., gene expression in alectinib-treated hepatoblasts vs. the average expression level across hepatoblasts treated with brigatinib, ceritinib, crizotinib, and lorlatinib). We aggregated single-cell counts into pseudobulk by individual, cell type, and treatment. For each cell type, we retained pseudobulk samples with at least 15 cells. We then performed voom-style normalization as implemented in dreamlet’s processAssays, using default min.count = 5, min.samples = 4, and min.prop = 0.4, and required at least four individuals per drug. There were 11 cell types did not pass these filters: acinar cells, adrenocortical cells, ductal cells, endocardial cells, ENS neurons, hematopoietic stem cells, islet endocrine cells, lens fibre cells, neuroendocrine cells, oligodendrocytes, and satellite cells. Since these filters were applied within each cell type, the set of tested genes is the same across all five 1v4 contrasts within a cell type, but can differ between cell types. The number of genes tested per cell type ranged from 4,527 (skeletal muscle cells) to 17,411 (radial glia), with a median of 10,226 genes.

To identify DEGs, we fit a linear mixed model with individual as a random effect. We applied multiple testing correction (Benjamini-Hochberg) within each 1v4 contrast across cell types. We define DEGs as genes with an adjusted p-value threshold < 0.05.

To estimate cell-type–specific effects across the 30 cell types passing our filters, we applied mashr to the dreamlet 1v4 coefficients using dreamlet::run_mash() separately per drug^40^. For genes not tested in a particular cell type, run_mash() sets the coefficient to zero and standard error to 1×10^6^ for the gene–cell-type pair before fitting the model, and then returns those posterior estimates as NA. We considered a gene to be a cell-type–specific DEG for a drug if it was tested in at least two cell types and was significant (lfsr < 0.05) in exactly one cell type and lfsr ≥0.05 in all others. We classified DEGs tested in only one cell type as singly-tested rather than cell-type–specific (Supplementary Data 8). To illustrate how DEGs may relate to adverse events, we examined DEGs in cell types relevant to selected adverse events, as described in Table 1.

To identify gene sets enriched among DEGs, we performed GSEA using fgseaMultilevel^41^ (nPermSimple=10,000, eps=0, seed=42). For each drug–cell-type combination, we ranked genes by the mash posterior mean divided by the standard deviation. We tested two MSigDB (*Homo sapiens* v.25.1.0) gene set collections: the Hallmark gene sets and GO BP terms^108–110^. We excluded gene sets with fewer than 15 or more than 500 genes. We performed multiple testing correction using the Benjamini-Hochberg method within each cell-type—treatment condition × gene set combination separately. We considered pathways with an adjusted p-value < 0.05 to be significant. Because mash shares information across cell types, we repeated GSEA as a sensitivity analysis with genes ranked by the pre-mash dreamlet t-statistic, using the same settings described above (Supplementary Data 10).

### Standard eQTL mapping

To prepare the filtered scRNA-seq data for eQTL mapping, we partitioned cells belonging to the same donor, cell type, and treatment into pseudobulk groups, requiring that each pseudobulk sample have at least 15 cells. We removed cell types present in fewer than 20 donors per treatment group, resulting in 21 cell types that passed filtering in both treatment and control conditions (the nine excluded cell types are: cardiomyocytes, erythroblasts, intestinal epithelial cells, lymphoid cells, megakaryocytes, myeloid cells, skeletal muscle cells, smooth muscle cells, and stellate cells). We chose this minimum donor cutoff in order to include cell types that are biologically relevant to *ALK* inhibitor toxicity (e.g., bronchiolar and alveolar epithelial cells and retinal pigment cells) while retaining sufficient statistical power to detect eQTLs. For each cell type, we performed trimmed mean of m-values (TMM) normalization on the scRNA-seq reads to obtain log(CPM) values. We then filtered genes to include those with a median UMI of at least 1 in addition to non-zero variance. We chose lenient filtering criteria in order to retain a sufficient number of genes for joint analysis across conditions in mash (described below). We log-transformed normalized reads for each gene using inverse normal transformation across samples.

To prepare YRI genotype data^100^ for eQTL mapping, we filtered out SNPs with a MAF less than 0.05. We also removed loci with rare genotypes within our sample (i.e., less than two homozygotes across samples in each cell-type—treatment condition). For each cell-type—treatment condition, we used MatrixEQTL to map *cis*-eQTLs, testing all variants within 100kb upstream and downstream of a gene’s TSS. For covariates, we used sex and expression-based principal components to capture hidden technical or biological factors of expression variability. We determined the number of expression-based PCs to include for each cell-type—treatment condition based on an elbow method using PCAforQTL^111^ (Supplementary Data 18). We computed the correlation of the selected PCs and other covariates and removed those highly correlated with any of the selected PCs (R^2^ > 0.9).

After mapping eQTLs separately in each condition, we used the *fastqtl2mash* package to reformat the MatrixEQTL results for mash^112^. Mash employs the lfsr to measure significance, which is more stringent than a false discovery rate because true discoveries are required to be not only nonzero but also correctly signed, incorporating the confidence in the direction of the effect. Posterior effect sizes are shrunk toward zero in proportion to their estimation uncertainty, so comparisons based on these estimates (i.e., our fold-change criterion used to define response eQTLs) are less sensitive to sampling noise than they would be using raw, per-condition regression coefficients. For each gene, the strong set used to learn covariance structure used the SNPs with the lowest p-value across all conditions. We ran mash jointly across all treatment and control conditions, testing genes that passed eQTL mapping filters in all conditions. We fit a single joint model (rather than fitting per cell type or per treatment) because we wanted the lfsr and effect sizes to be comparable across both treatment and cell type axes. Given the large number of conditions tested, we estimated null correlation structure among conditions using the expectation-maximization algorithm implemented in mashr^113–115^.

### Identifying response eQTLs using mash

Gene-SNP pairs passing the significance threshold of lfsr < 0.05 are considered eQTLs. To identify response eQTLs, we asked whether eQTL effects changed between a given cell type in the focal treatment group vs. the control group. We defined response eQTLs separately for each cell type. For a significant SNP-gene pair to be considered a response eQTL, it needed to be significant (lfsr < 0.05) in either the treatment or control group. We also required response eQTL effect sizes to differ either in their magnitude (at least a three-fold difference between conditions) or in the direction of their effects.

### Genotype-by-treatment interaction model

To evaluate our mash-based definition of response eQTLs against a test of the genotype-by-treatment difference, we refit each tested gene-SNP pair using an interaction model in MatrixEQTL (modelLINEAR_CROSS). For each cell type, we combined pseudobulk expression for a given drug and tested a genotype x treatment interaction term with sex and expression PCs (per cell type) as covariates. Genes and SNPs were restricted to those tested in mash so that the estimates were comparable. We applied eigenMT^116^ correction for linkage disequilibrium among tested SNPs per gene and added a secondary correction across genes within each drug using Benjamini-Hochberg. We compared the interaction coefficient with the mash posterior effect-size difference (Δβ = β_treatment_ – β_control_) using Pearson’s correlation coefficient and the proportion of pairs with a concordant sign.

### Enrichment of eQTLs and eGenes in regulatory elements and gene-level features

To assess enrichment of eQTLs within annotated regulatory elements, we obtained candidate *cis* regulatory elements (cCREs) from ENCODE v4, using their definitions of promoter-like signatures (PLS), proximal enhancer like signatures (pELS), and distal enhancer like signatures (dELS)^117^. PLSs are defined as being within 200bp (center to center) of an annotated GENCODE TSS or experimentally derived TSS, along with having high chromatin accessibility and H3K4me3 signals. ELSs are defined as having high DNase and H3K27ac, and need to have low H3K4me3 if they are within 200 bp of an annotated TSS. ELS within 2kb of a TSS are considered proximal and all others are considered distal. We ran Fisher’s exact test to directly compare enrichment of response eQTLs against that of non-response eQTLs in each cCRE class.

We then asked whether certain gene-level features are enriched among response eGenes and non-response eGenes. We focused on three metrics, including probability of loss-of-function intolerance (pLI; annotated from gnomad v4.1)^118^, TSS counts calculated from FANTOM data^119^, and activity-by-contact (ABC) cumulative enhancer length^120,121^, which predicts enhancer-gene links using chromatin accessibility, H3K27ac, Hi-C 3D contact data. For each gene, we summed enhancer lengths across all active cell types and averaged across cell types. For each condition, we calculated these three features for response and non-response eGenes and used a one-sided sign test to assess how consistently (across all conditions) each feature was higher in response eGenes.

### Mapping DEGs and response eQTLs to adverse events

To connect response eQTLs to the adverse events associated with each drug, we annotated each response eQTL with adverse events listed for that drug in Section 5 (Warnings and Precautions) of its FDA label (Table 1). We matched a response eQTL to an adverse event when the cell type in which it was identified is one of the HDC cell types plausibly relevant to the affected tissue; therefore, a response eQTL can be annotated with more than one adverse event. We considered all adverse events listed for a drug rather than restricting only to the most frequent or most characteristic ones because inter-individual genetic variation could also contribute to adverse events that are relatively less frequent for a drug. Adverse events that overlap between drugs, such as hepatotoxicity, could also have different mechanisms in different drugs and could potentially be affected differently by genetic variation, so these were annotated for each drug rather than only as a single class-wide event.

## Acknowledgements

We thank Wenhe Lin, Ben Umans, and Erik McIntire for discussions. This work was supported by the Cancer Research Foundation as well as R35GM131726 to YG. E.G.W. was supported by F30HD118747 and T32GM150375.

## Author contributions

Conceptualization, Y.G.

Methodology, E.G.W., K.L.R., K.A.B., and Y.G.

Formal analysis, E.G.W.

Funding acquisition, E.E.V and Y.G.

Supervision, Y.G.

Data collection, K.L.R., E.M.

Writing: original draft, E.G.W.

Writing: review and editing, E.G.W., K.L.R., N.M.G., K.A.B, E.M., E.E.V., and Y.G.

## Competing interests

K.L.R. and Y.G. are co-founders and equity holders of CellCipher, Inc. K.L.R. and Y.G. are co-inventors on patent application 18067192 related to the *in vitro* model used in this work

## Data availability

The single-cell RNA-seq data have been deposited at GEO under accession GSE347774.

Genotype data for the YRI population is available through the International Genome Sample Resource at ISGR: https://www.internationalgenome.org/data-portal/population/YRI.

An interactive viewer for the supplementary results is available at: https://wooeg.shinyapps.io/alk-resource/

## Code availability

Code used to perform the analyses is available on Github: https://github.com/elizabethgeenaw/ALK

## Supplementary Files

Supplementary Figures.pdf

Supplementary Data.xlsx

## Notes

https://wooeg.shinyapps.io/alk-resource/

