## Supplemental Figures for "Genetic regulation of ALK inhibitor response in diverse human cell types"

### Supplementary Figures Legend

**Supplementary Fig. 1.** Cell type composition by individual and treatment
**Supplementary Fig. 2**. DEG counts vs cell type abundance by drug
**Supplementary Fig. 3.** Volcano plots of 1v4 DE for each cell type and ALK inhibitor
**Supplementary Fig. 4**. Number of cell-type–specific DEGs per drug
**Supplementary Fig. 5.** Hallmark gene set enrichment (NES) across ALK inhibitors and cell types
**Supplementary Fig. 6**. GO BP gene set enrichment (NES) for each ALK inhibitor, pre-mash
**Supplementary Fig. 7**. GO BP pathway enrichment (NES) for each ALK inhibitor, post-mash
**Supplementary Fig. 8**. Number of significant cis-eQTLs per cell-type–treatment condition
**Supplementary Fig. 9**. Standard cis-eQTL counts vs pseudobulk donor and cell numbers
**Supplementary Fig. 10**. Concordance between mash response eQTL effect sizes and a genotype-treatment interaction model
**Supplementary Fig. 11**. SNP-to-TSS distance for response and non-response eQTLs
**Supplementary Fig. 12**. cCRE class enrichment (PLS, pELS, dELS) for response vs non-response eQTLs

### Supplementary Data Legend

**Supplementary Data 1.** Metadata for each treatment library
**Supplementary Data 2.** Per-sample QC and cell-type summary of the treatment data
**Supplementary Data 3.** Per-cell type cell counts and frequencies in treatment and control
**Supplementary Data 4**. Per-donor cell counts and QC metrics for the treatment and control
**Supplementary Data 5.** Significant 1v4 DEGs
**Supplementary Data 6.** Number of cell types in which a DEG was significant, before mash
**Supplementary Data 7.** Number of cell types in which a DEG was significant, after mash
**Supplementary Data 8.** Cell-type-specific DEGs
**Supplementary Data 9.** Differential expression GSEA, post-mash
**Supplementary Data 10**. Differential expression GSEA, pre-mash
**Supplementary Data 11.** All significant cis-eQTLs
**Supplementary Data 12.** All response eQTLs
**Supplementary Data 13.** cCRE enrichment of response vs non-response eQTLs
**Supplementary Data 14.** Fold-change sensitivity analysis of gene-level features
**Supplementary Data 15.** Response eQTLs anchored to Table 1 (Warnings and Precautions) of FDA label
**Supplementary Data 16.** Drug-specific response eQTLs within a cell type
**Supplementary Data 17**. Condition (treatment and cell-type)-specific response eQTLs
**Supplementary Data 18**. Expression PC covariates per condition

**Supplementary Figure 1.** Cell type composition by individual and treatment.

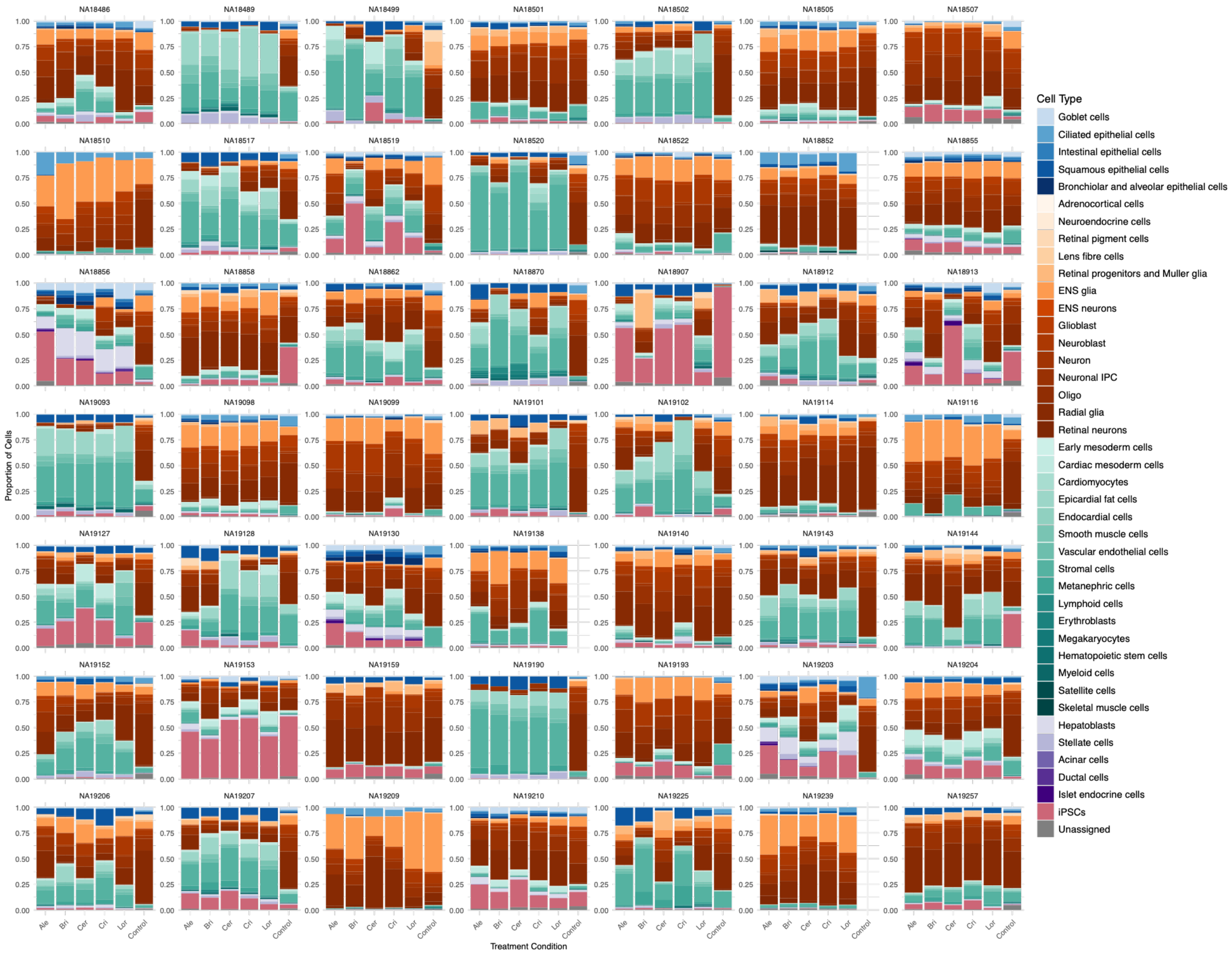

**Supplementary Figure 2.** Scatter plot of DEG count (y-axis, log10 scale; FDR < 0.05) and cell type abundance summed across individuals (x-axis, log10 scale). Each dot represents a cell type. Each panel is a drug and Spearman correlations for each drug are reported. The dashed line and shaded band show an OLS fit with 95% confidence interval.

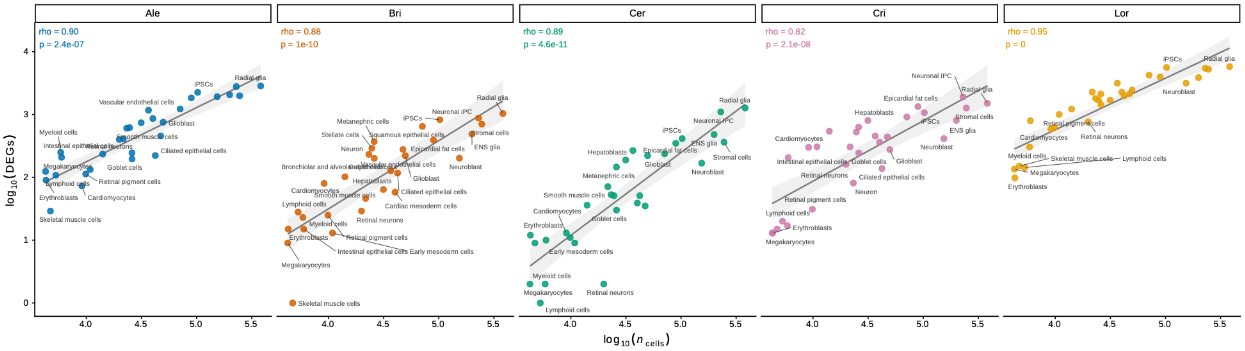

**Supplementary Figure 3.** Volcano plots for each cell type and 1v4 DE comparison. Each panel shows a cell type (row) and five ALK inhibitors (columns). Each point represents one gene. The x-axis is log2 fold change of one treatment relative to the mean of the other four, the y-axis is -log10(p). The dashed horizontal line marks the p-value corresponding to FDR = 0.05 (Benjamini-Hochberg). Colored points indicate DEGs at FDR < 0.05. The top six genes (by p-value) per panel are labeled. The colored vertical band on the left identifies the cell type and band color indicates germ layer: epithelium (blue), ectoderm (orange), mesoderm (teal), endoderm (purple), iPSCs (pink).

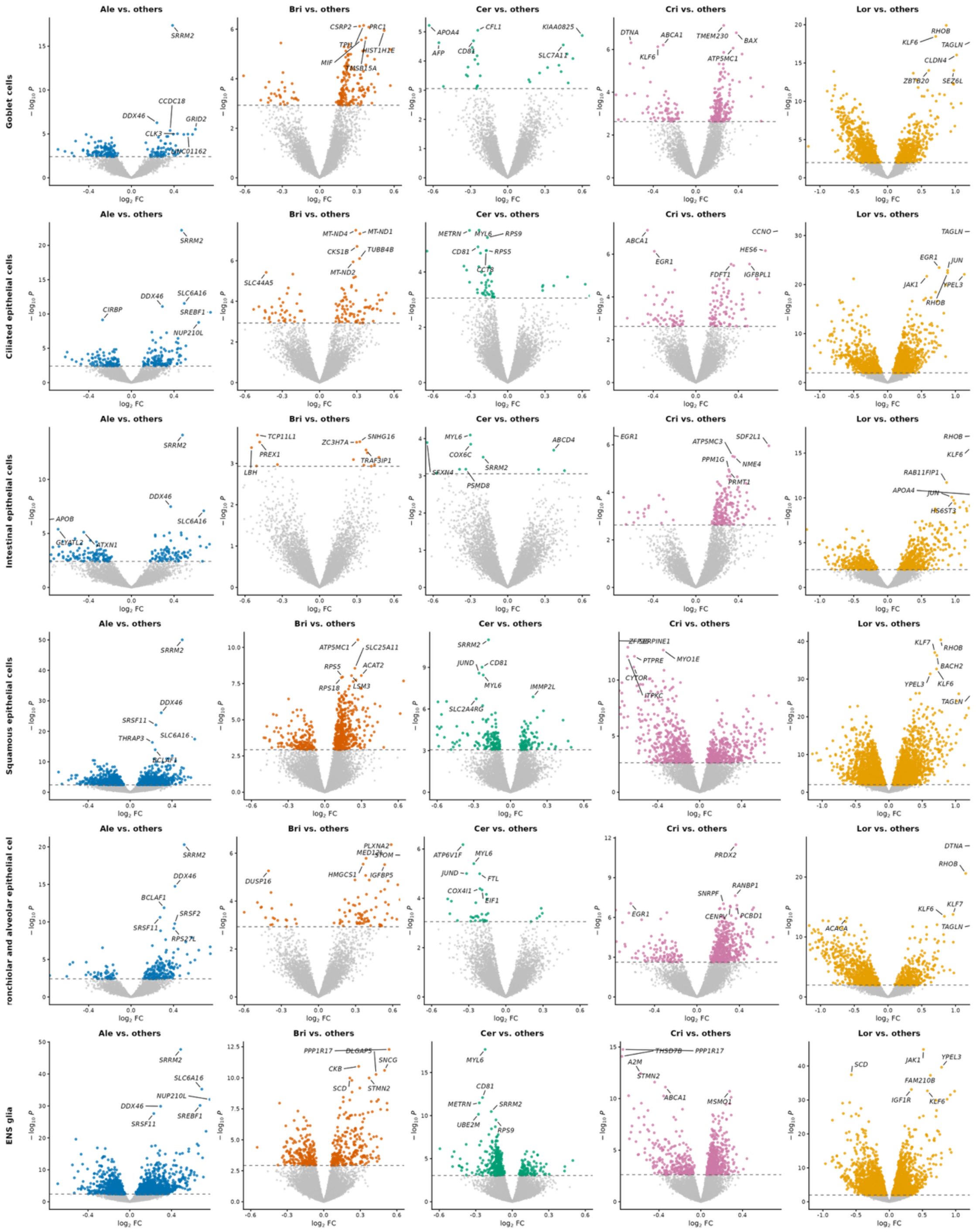

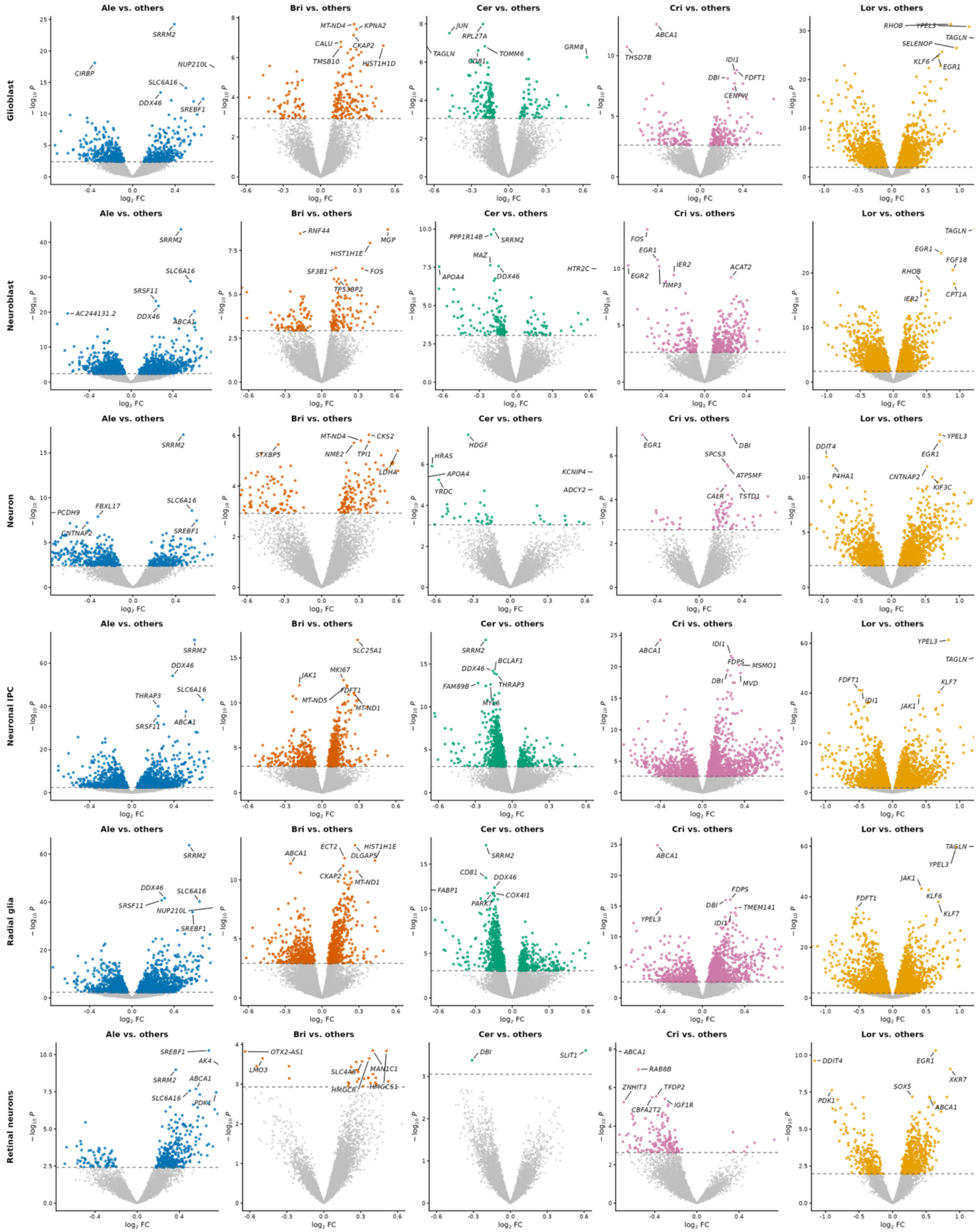

Ale vs. others
Bri vs. others
Cer vs. others
Cri vs. others
Lor vs. others
Glioblast
Neuroblast
Neuron
Neuronal IPC
Radial glia
Retinal neurons
$-\log_{10} P$
$\log_2$ FC
SRRM2
CIRBP
NUP210L
SLC6A16
DDX46
SREBF1
MT-ND4
KPNA2
CALU
CKAP2
TMSB10
HIST1H1D
JUN
RPL27A
TAGLN
TOMM6
GRM8
CD81
ABCA1
THSD7B
IDI1
DBI
FDFT1
CENPW
RHOB
YPEL3
SELENOP
KLF6
EGR1
AC244131.2
SRSF11
RNF44
MGP
HIST1H1E
SF3B1
FOS
TP53BP2
PPP1R14B
MAZ
HTR2C
APOA4
EGR2
IER2
TIMP3
ACAT2
FGF18
CPT1A
PCDH9
FBXL17
CNTNAP2
STXBP5
CKS2
NME2
TPI1
LDHA
HDGF
HRAS
KCNIP4
ADCY2
YRDC
SPCS3
ATP5MF
CALR
TSTD1
DDIT4
P4HA1
KIF3C
THRAP3
SLC25A1
JAK1
MKI67
MT-ND5
MT-ND1
BCLAF1
FAM89B
MYL6
FDPS
MSMO1
MVD
KLF7
ECT2
DLGAP5
FABP1
COX4I1
PARK7
TMEM141
AK4
PDK1
OTX2-AS1
LMO3
SLC4A8
MAN1C1
HMGCR
HMGCS1
SLIT1
RAB8B
ZNHIT3
TFDP2
CBFA2T2
IGF1R
XKR7
SOX5

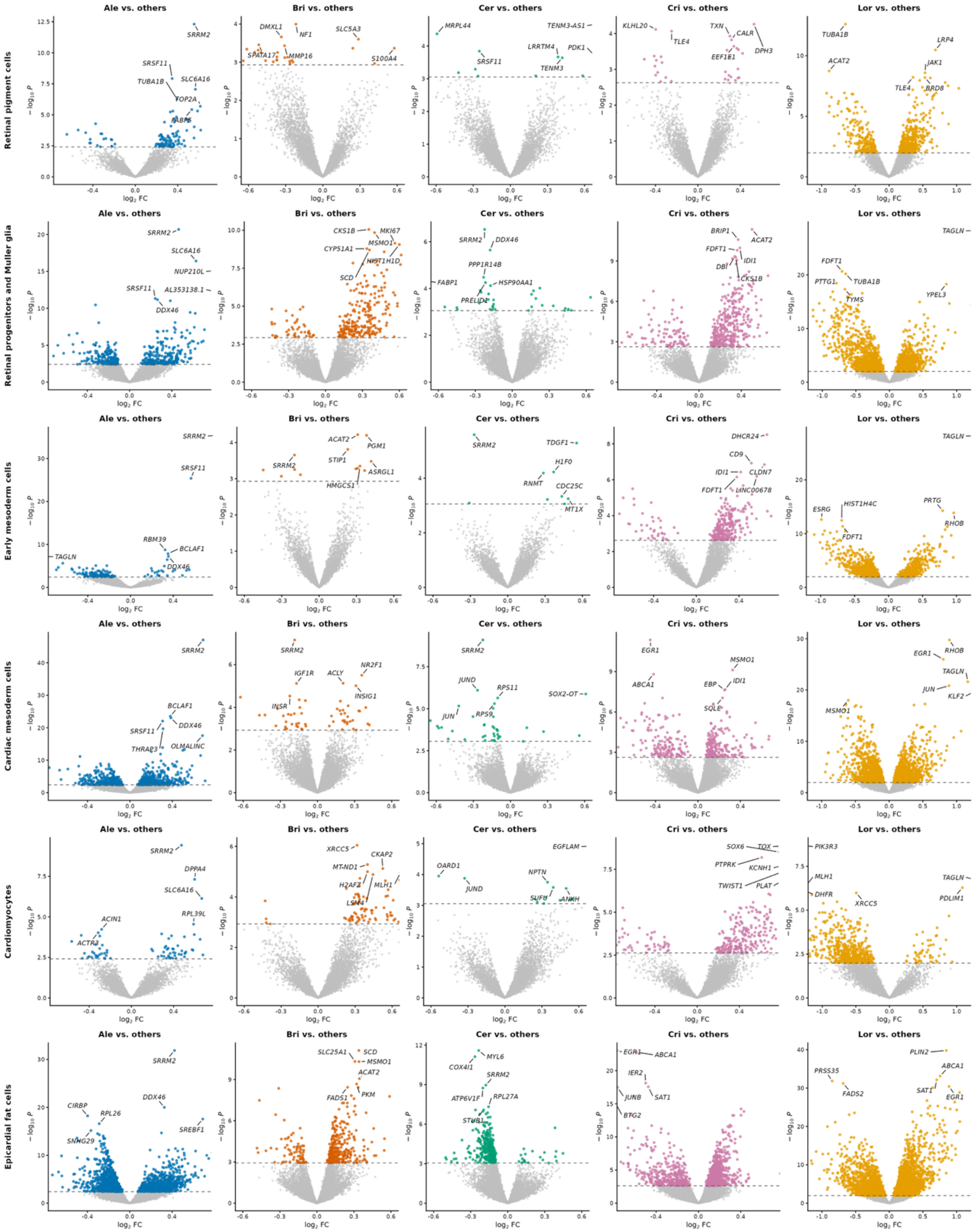

Ale vs. others
Bri vs. others
Cer vs. others
Cri vs. others
Lor vs. others
Retinal pigment cells
Retinal progenitors and Muller glia
Early mesoderm cells
Cardiac mesoderm cells
Cardiomyocytes
Epicardial fat cells
log2 FC
− log10 P
SRRM2
SRSF11
TUBA1B
SLC6A16
TOP2A
FABP5
DMXL1
NF1
SLC5A3
SPATA17
MMP16
S100A4
MRPL44
TENM3-AS1
LRRTM4
PDK1
TENM3
KLHL20
TLE4
TXN
CALR
EEF1E1
DPH3
ACAT2
LRP4
JAK1
BRD8
NUP210L
AL353138.1
DDX46
CKS1B
MKI67
MSMO1
CYP51A1
HIST1H1D
SCD
PPP1R14B
FABP1
HSP90AA1
PRELID1
BRIP1
FDFT1
IDI1
DBI
TAGLN
PTTG1
TYMS
YPEL3
RBM39
BCLAF1
PGM1
STIP1
ASRGL1
HMGCS1
TDGF1
H1F0
RNMT
CDC25C
MT1X
DHCR24
CD9
CLDN7
LINC00678
ESRG
HIST1H4C
PRTG
RHOB
THRAP3
OLMALINC
IGF1R
ACLY
NR2F1
INSR
INSIG1
JUND
RPS11
SOX2-OT
JUN
RPS9
EGR1
ABCA1
EBP
SQLE
KLF2
DPPA4
ACIN1
RPL39L
ACTR3
XRCC5
CKAP2
MT-ND1
H2AFZ
MLH1
LSM4
EGFLAM
OARD1
NPTN
SUFU
ANKH
SOX6
TOX
PTPRK
KCNH1
TWIST1
PLAT
PIK3R3
DHFR
PDLIM1
CIRBP
RPL26
SNHG29
SREBF1
SLC25A1
FADS1
PKM
MYL6
COX4I1
ATP6V1F
RPL27A
STUB1
IER2
SAT1
JUNB
BTG2
PLIN2
PRSS35
FADS2

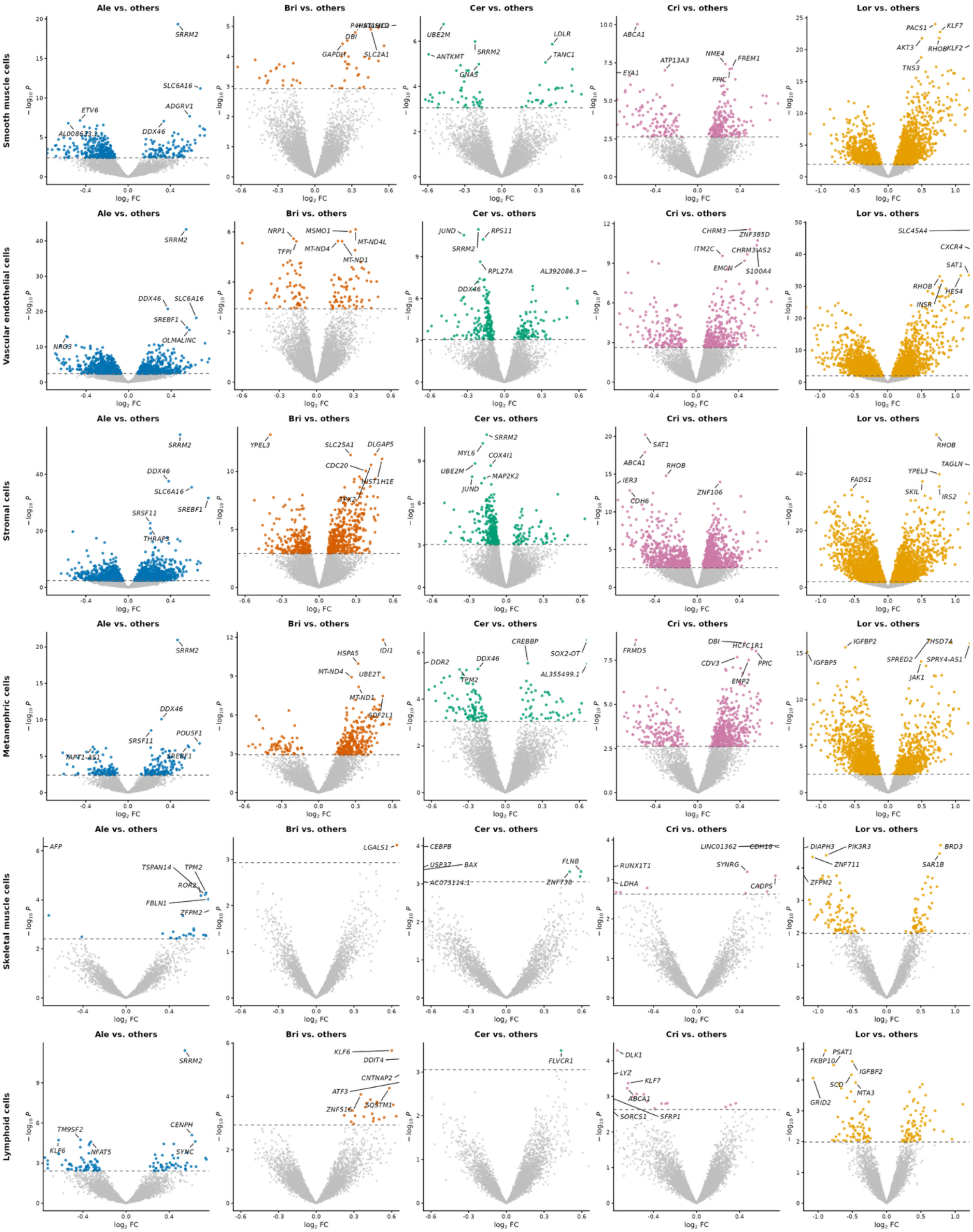

Smooth muscle cells
Vascular endothelial cells
Stromal cells
Metanephric cells
Skeletal muscle cells
Lymphoid cells
Ale vs. others
Bri vs. others
Cer vs. others
Cri vs. others
Lor vs. others
$-\log_{10} P$
$\log_2$ FC
SRRM2
SLC6A16
ADGRV1
DDX46
ETV6
DBI
GAPDH
SLC2A1
UBE2M
ANTKMT
GNAS
LDLR
TANC1
ABCA1
ATP13A3
NME4
FREM1
PPIC
EYA1
PACS1
KLF7
AKT3
RHOB
KLF2
TNS3
SREBF1
OLMALINC
NRG3
NRP1
MSMO1
TFPI
MT-ND4
MT-ND4L
MT-ND1
JUND
RPS11
RPL27A
AL392086.3
CHRM3
ZNF385D
ITM2C
CHRM3-AS2
EMGN
S100A4
SLC45A4
CXCR4
SAT1
HES4
INSR
SRSF11
THRAP3
YPEL3
SLC25A1
DLGAP5
CDC20
HIST1H1E
MYL6
COX4I1
MAP2K2
IER3
CDH6
ZNF106
FADS1
TAGLN
SKIL
IRS2
POU5F1
IDI1
HSPA5
UBE2T
SDF2L1
CREBBP
SOX2-OT
DDR2
TPM2
AL355499.1
FRMD5
HCFC1R1
CDV3
EMP2
IGFBP2
THSD7A
IGFBP5
SPRED2
SPRY4-AS1
JAK1
AFP
TSPAN14
ROR2
FBLN1
ZFPM2
LGALS1
CEBPB
USP37
BAX
AC073114.1
FLNB
ZNF738
RUNX1T1
LDHA
LINC01362
CDH18
SYNGR
CADPS
DIAPH3
PIK3R3
ZNF711
BRD3
SAR1B
TM9SF2
KLF6
NFAT5
CENPH
SYNC
DDIT4
CNTNAP2
ATF3
ZNF516
SQSTM1
FLVCR1
DLK1
LYZ
SORCS1
SFRP1
PSAT1
FKBP10
SCD
MTA3
GRID2

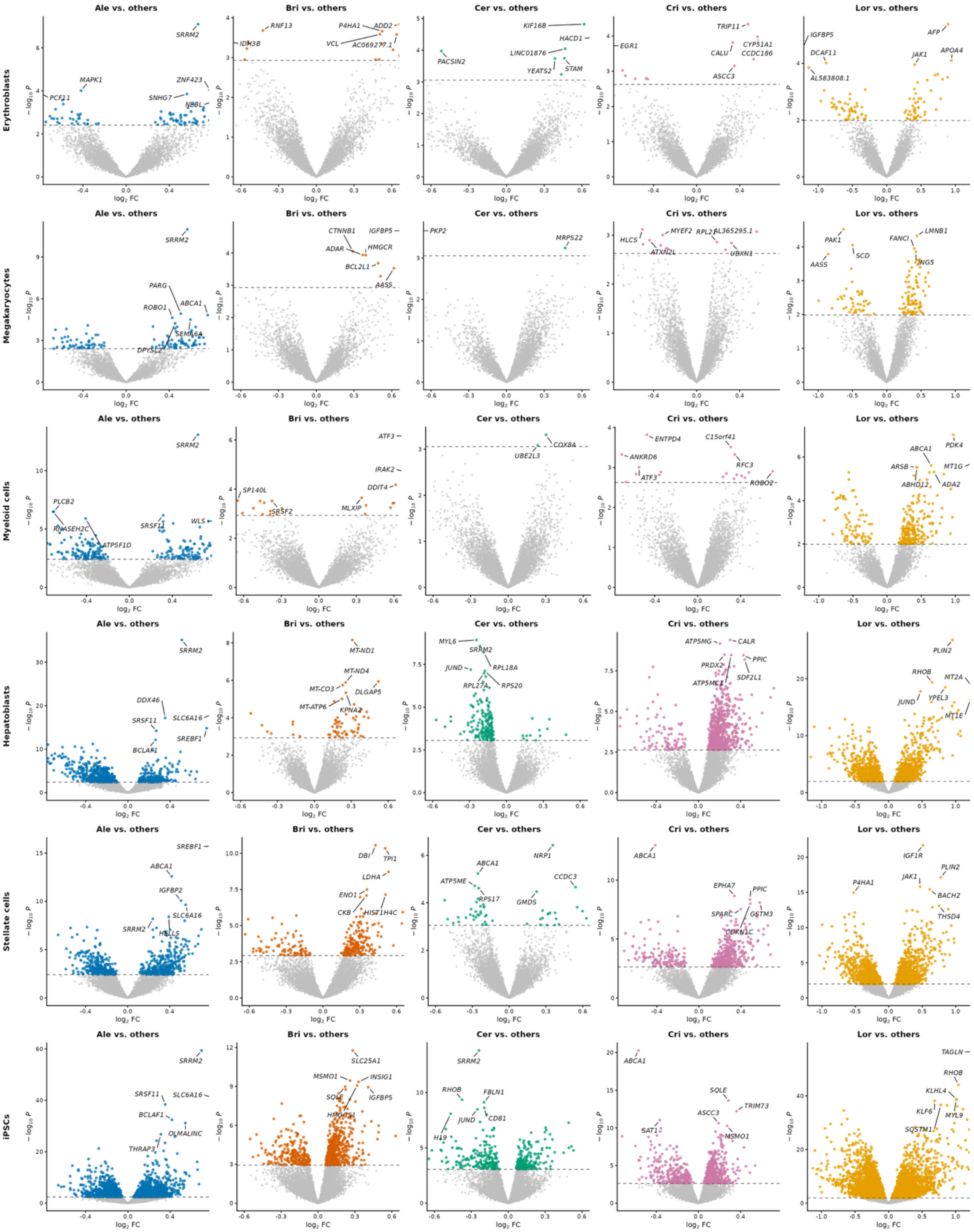

Ale vs. others
Bri vs. others
Cer vs. others
Cri vs. others
Lor vs. others
Erythroblasts
Megakaryocytes
Myeloid cells
Hepatoblasts
Stellate cells
iPSCs
log2 FC
−log10 P
SRRM2
MAPK1
PCF11
ZNF423
SNHG7
NBBL
RNF13
IDH3B
P4HA1
ADD2
VCL
AC069277.1
KIF16B
HACD1
PACSIN2
LINC01876
YEATS2
STAM
EGR1
TRIP11
CYP51A1
CALU
CCDC186
ASCC3
IGFBP5
DCAF11
AL583808.1
AFP
APOA4
JAK1
PARG
ROBO1
ABCA1
SEMA6A
DPYSL2
CTNNB1
ADAR
HMGCR
BCL2L1
AASS
PKP2
MRPS22
HLCS
MYEF2
RPL22
AL365295.1
ATXN2L
UBXN1
PAK1
SCD
FANCI
LMNB1
ING5
PLCB2
RNASEH2C
ATP5F1D
SRSF11
WLS
ATF3
IRAK2
DDIT4
SP140L
SRSF2
MLXIP
COX8A
UBE2L3
ENTPD4
C15orf41
ANKRD6
RFC3
ROBO2
PDK4
ARSB
MT1G
ABHD12
ADA2
DDX46
SLC6A16
SREBF1
BCLAF1
MT-ND1
MT-ND4
MT-CO3
DLGAP5
MT-ATP6
KPNA2
MYL6
JUND
RPL18A
RPL27A
RPS20
ATP5MG
CALR
PRDX2
PPIC
ATP5MC1
SDF2L1
PLIN2
RHOB
MT2A
YPEL3
MT1E
IGFBP2
HELLS
DBI
TPI1
LDHA
ENO1
CKB
HIST1H4C
NRP1
ATP5ME
CCDC3
RPS17
GMDS
EPHA7
SPARC
GSTM3
CDKN1C
IGF1R
P4HA1
BACH2
THSD4
OLMALINC
THRAP3
SLC25A1
MSMO1
INSIG1
SQLE
HMGCS1
RHOB
FBLN1
CD81
H19
TRIM73
SAT1
TAGLN
KLHL4
KLF6
MYL9
SQSTM1

**Supplementary Figure 4.** Number of cell-type–specific DEGs and singly tested genes. Singly tested genes were genes tested only in a single cell type, while cell-type–specific were tested in at least two.

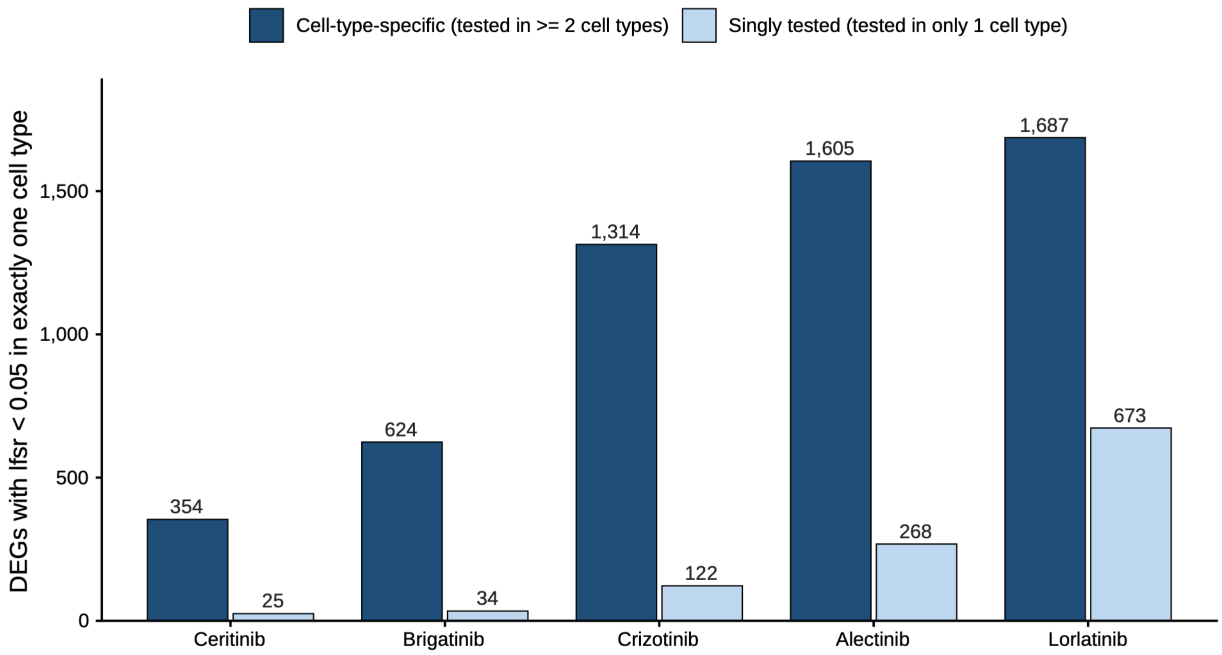

**Supplementary Figure 5.** Hallmark pathway enrichment across ALK inhibitors from GSEA using (A) pre-mash dreamlet t-statistics and (B) post-mash statistics (posterior mean / SD). Each cell shows the Normalized Enrichment Score (NES) for a Hallmark gene set for each drug–cell-type combination. Blue indicates higher expression relative to the other four ALK inhibitors; orange indicates lower expression. Each row is a cell type, grouped by germ layer, and each column is a pathway. * indicates significant enrichment (p-adj < 0.05) for that cell-type–drug.

**A**

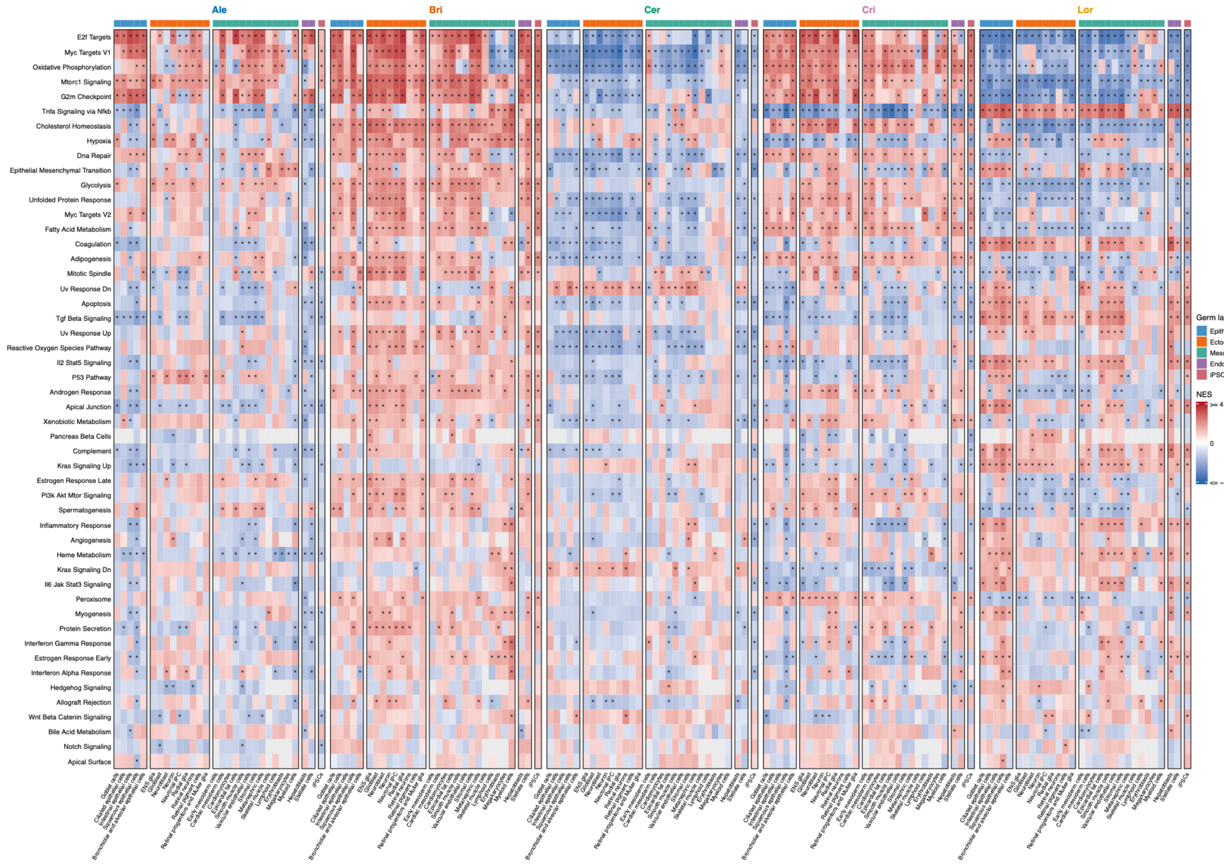

B

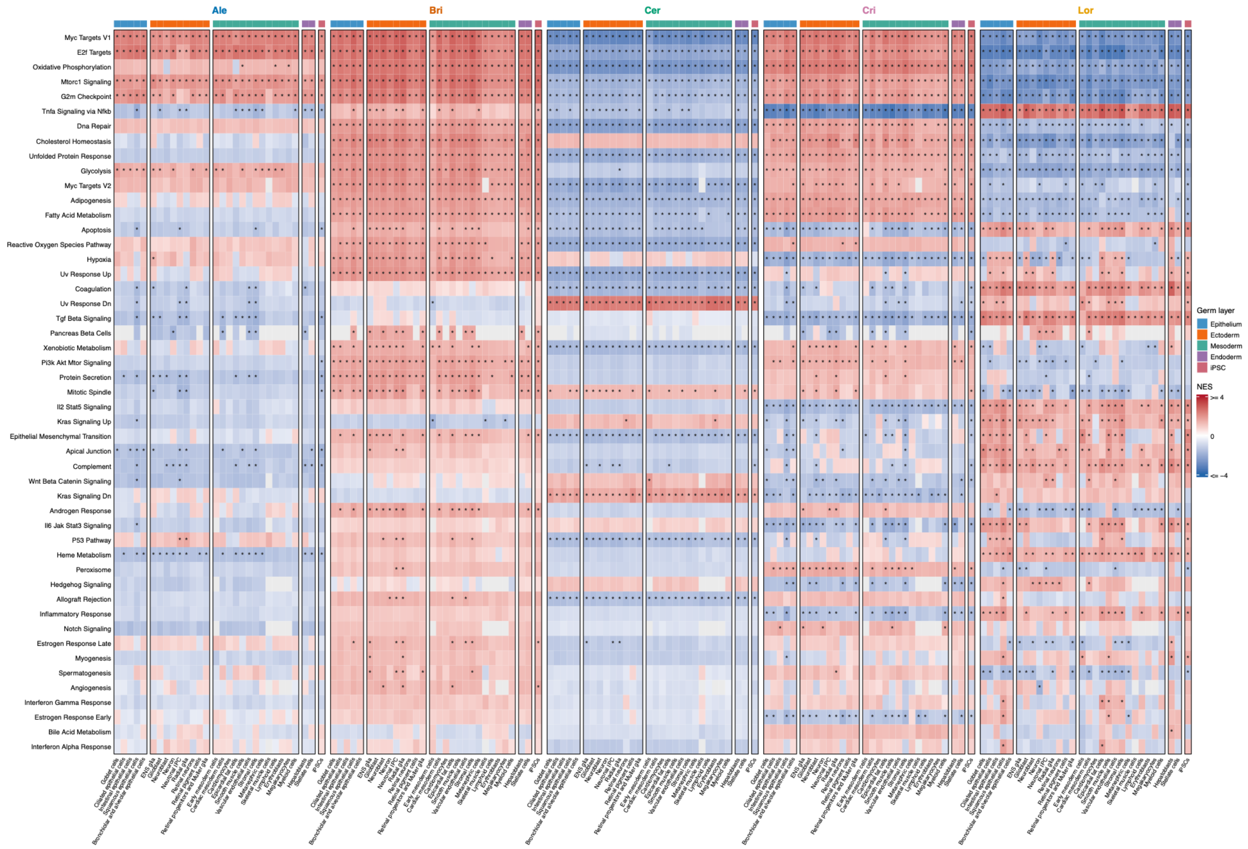
Ale
Bri
Cer
Cri
Lor
Myc Targets V1
E2f Targets
Oxidative Phosphorylation
Mtorc1 Signaling
G2m Checkpoint
Tnfa Signaling via Nfkb
Dna Repair
Cholesterol Homeostasis
Unfolded Protein Response
Glycolysis
Myc Targets V2
Adipogenesis
Fatty Acid Metabolism
Apoptosis
Reactive Oxygen Species Pathway
Hypoxia
Uv Response Up
Coagulation
Uv Response Dn
Tgf Beta Signaling
Pancreas Beta Cells
Xenobiotic Metabolism
Pi3k Akt Mtor Signaling
Protein Secretion
Mitotic Spindle
Il2 Stat5 Signaling
Kras Signaling Up
Epithelial Mesenchymal Transition
Apical Junction
Complement
Wnt Beta Catenin Signaling
Kras Signaling Dn
Androgen Response
Il6 Jak Stat3 Signaling
P53 Pathway
Heme Metabolism
Peroxisome
Hedgehog Signaling
Allograft Rejection
Inflammatory Response
Notch Signaling
Estrogen Response Late
Myogenesis
Spermatogenesis
Angiogenesis
Interferon Gamma Response
Estrogen Response Early
Bile Acid Metabolism
Interferon Alpha Response
Germ layer
Epithelium
Ectoderm
Mesoderm
Endoderm
iPSC
NES
>= 4
0
<= −4

**Supplementary Figure 6.** GO BP pathway enrichment for each ALK inhibitor from GSEA using pre-mash dreamlet computed t-statistics. Each cell of the heatmap shows the Normalized Enrichment Score (NES) for a GO BP gene set for each drug–cell-type combination. Blue indicates upregulation relative to the other four ALK inhibitors; orange indicates downregulation. The top 15 positive and negative NES pathways (i.e. largest max(NES) and most negative min(NES) across cell types with at least one significant cell type in that direction) are shown. Within each up/down block, pathways are ordered by their signed extreme NES. Each row is a cell type and each column is a pathway. * indicates significant enrichment (p-adj < 0.05). Gray boxes indicate cell-type–pathway combinations where NES could not be computed (for example, too few genes overlapped the gene set in that cell type).

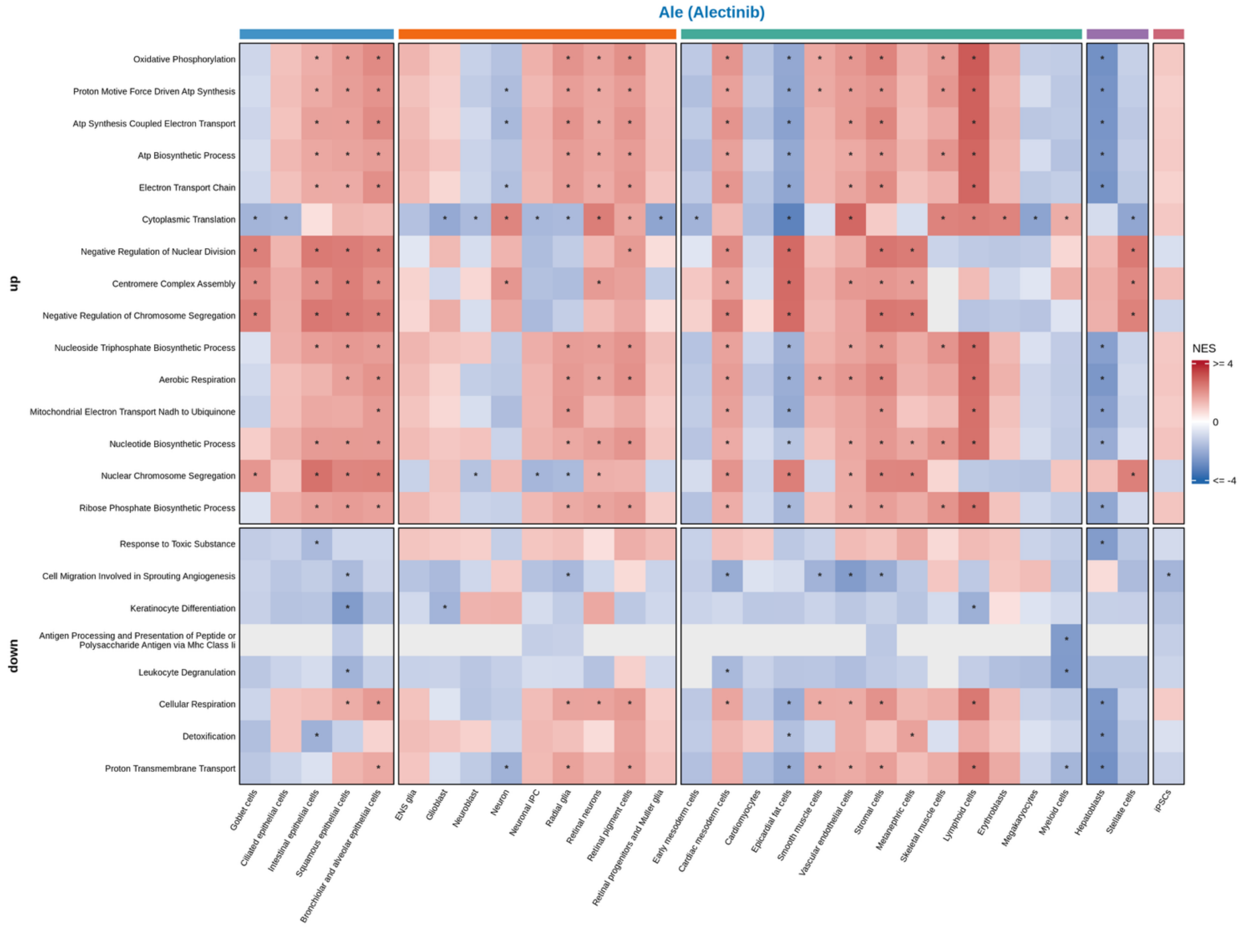

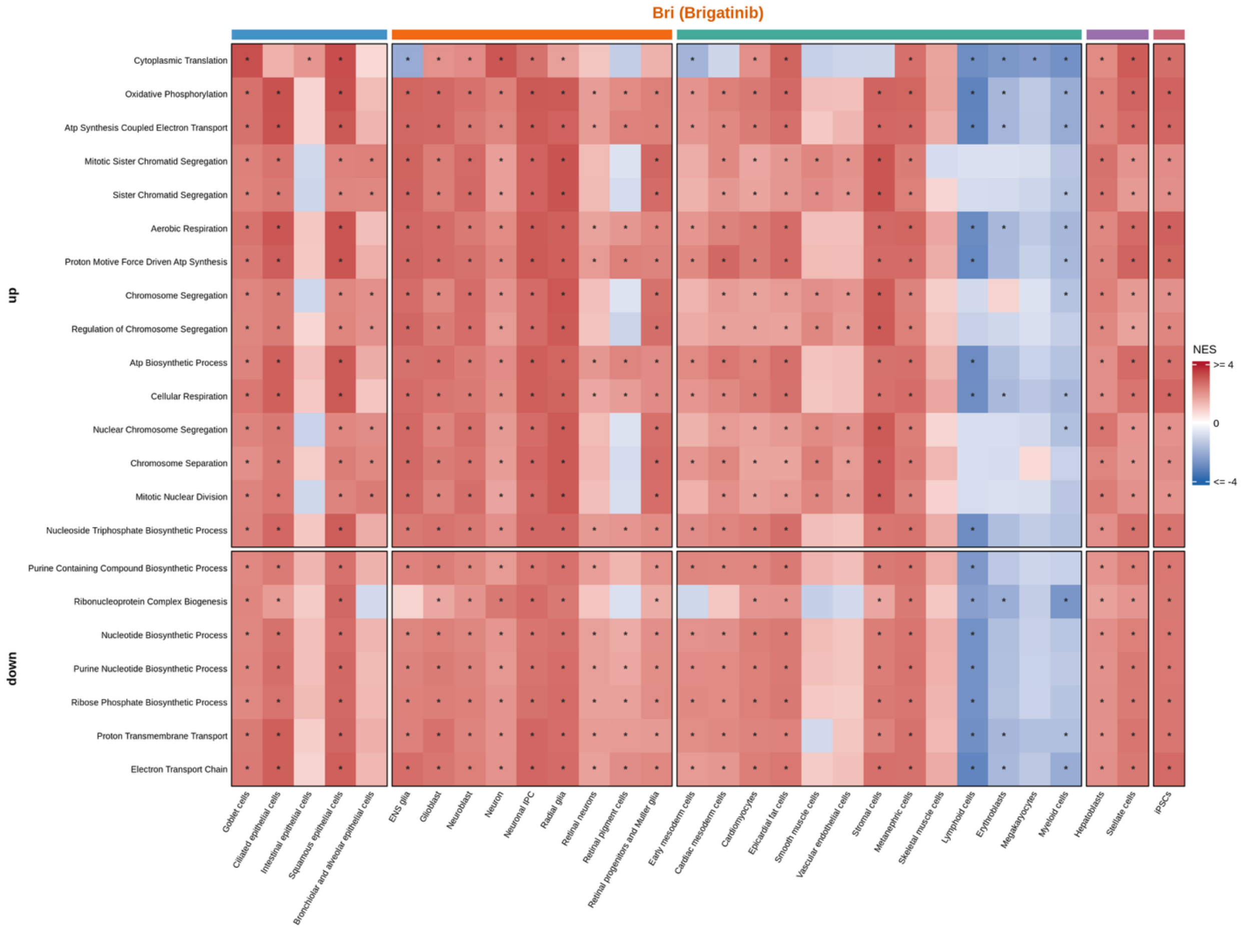
Bri (Brigatinib)
up
down
Cytoplasmic Translation
Oxidative Phosphorylation
Atp Synthesis Coupled Electron Transport
Mitotic Sister Chromatid Segregation
Sister Chromatid Segregation
Aerobic Respiration
Proton Motive Force Driven Atp Synthesis
Chromosome Segregation
Regulation of Chromosome Segregation
Atp Biosynthetic Process
Cellular Respiration
Nuclear Chromosome Segregation
Chromosome Separation
Mitotic Nuclear Division
Nucleoside Triphosphate Biosynthetic Process
Purine Containing Compound Biosynthetic Process
Ribonucleoprotein Complex Biogenesis
Nucleotide Biosynthetic Process
Purine Nucleotide Biosynthetic Process
Ribose Phosphate Biosynthetic Process
Proton Transmembrane Transport
Electron Transport Chain
Goblet cells
Ciliated epithelial cells
Intestinal epithelial cells
Squamous epithelial cells
Bronchiolar and alveolar epithelial cells
ENS glia
Glioblast
Neuroblast
Neuron
Neuronal IPC
Radial glia
Retinal neurons
Retinal pigment cells
Retinal progenitors and Muller glia
Early mesoderm cells
Cardiac mesoderm cells
Cardiomyocytes
Epicardial fat cells
Smooth muscle cells
Vascular endothelial cells
Stromal cells
Metanephric cells
Skeletal muscle cells
Lymphoid cells
Erythroblasts
Megakaryocytes
Myeloid cells
Hepatoblasts
Stellate cells
iPSCs
NES
>= 4
0
<= -4

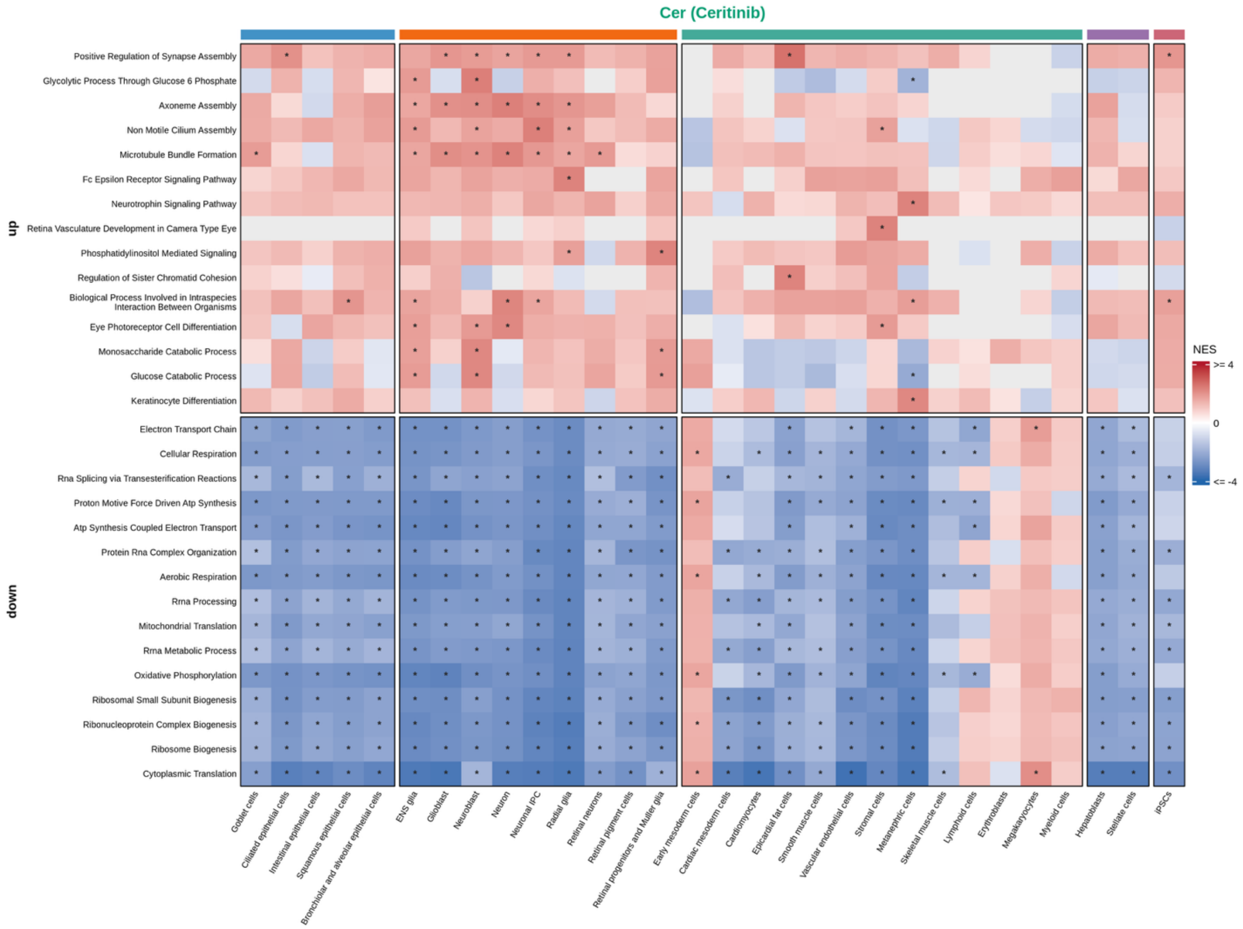
Cer (Ceritinib)
up
down
Positive Regulation of Synapse Assembly
Glycolytic Process Through Glucose 6 Phosphate
Axoneme Assembly
Non Motile Cilium Assembly
Microtubule Bundle Formation
Fc Epsilon Receptor Signaling Pathway
Neurotrophin Signaling Pathway
Retina Vasculature Development in Camera Type Eye
Phosphatidylinositol Mediated Signaling
Regulation of Sister Chromatid Cohesion
Biological Process Involved in Intraspecies Interaction Between Organisms
Eye Photoreceptor Cell Differentiation
Monosaccharide Catabolic Process
Glucose Catabolic Process
Keratinocyte Differentiation
Electron Transport Chain
Cellular Respiration
Rna Splicing via Transesterification Reactions
Proton Motive Force Driven Atp Synthesis
Atp Synthesis Coupled Electron Transport
Protein Rna Complex Organization
Aerobic Respiration
Rrna Processing
Mitochondrial Translation
Rrna Metabolic Process
Oxidative Phosphorylation
Ribosomal Small Subunit Biogenesis
Ribonucleoprotein Complex Biogenesis
Ribosome Biogenesis
Cytoplasmic Translation
Goblet cells
Ciliated epithelial cells
Intestinal epithelial cells
Squamous epithelial cells
Bronchiolar and alveolar epithelial cells
ENS glia
Glioblast
Neuroblast
Neuron
Neuronal IPC
Radial glia
Retinal neurons
Retinal pigment cells
Retinal progenitors and Muller glia
Early mesoderm cells
Cardiac mesoderm cells
Cardiomyocytes
Epicardial fat cells
Smooth muscle cells
Vascular endothelial cells
Stromal cells
Metanephric cells
Skeletal muscle cells
Lymphoid cells
Erythroblasts
Megakaryocytes
Myeloid cells
Hepatoblasts
Stellate cells
iPSCs
NES
>= 4
0
<= -4

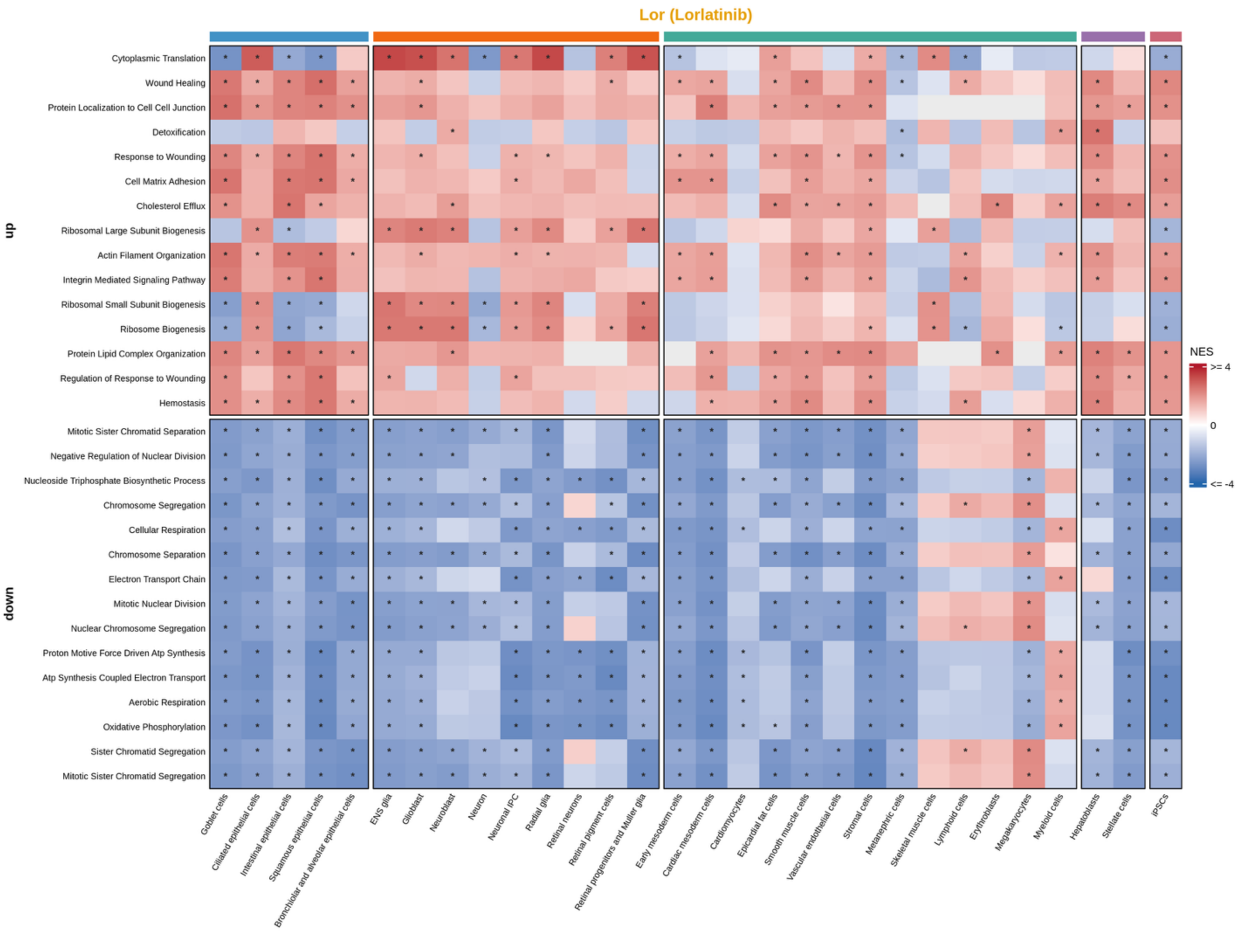

Lor (Lorlatinib)
up
down
Cytoplasmic Translation
Wound Healing
Protein Localization to Cell Cell Junction
Detoxification
Response to Wounding
Cell Matrix Adhesion
Cholesterol Efflux
Ribosomal Large Subunit Biogenesis
Actin Filament Organization
Integrin Mediated Signaling Pathway
Ribosomal Small Subunit Biogenesis
Ribosome Biogenesis
Protein Lipid Complex Organization
Regulation of Response to Wounding
Hemostasis
Mitotic Sister Chromatid Separation
Negative Regulation of Nuclear Division
Nucleoside Triphosphate Biosynthetic Process
Chromosome Segregation
Cellular Respiration
Chromosome Separation
Electron Transport Chain
Mitotic Nuclear Division
Nuclear Chromosome Segregation
Proton Motive Force Driven Atp Synthesis
Atp Synthesis Coupled Electron Transport
Aerobic Respiration
Oxidative Phosphorylation
Sister Chromatid Segregation
Mitotic Sister Chromatid Segregation
Goblet cells
Ciliated epithelial cells
Intestinal epithelial cells
Squamous epithelial cells
Bronchiolar and alveolar epithelial cells
ENS glia
Glioblast
Neuroblast
Neuron
Neuronal IPC
Radial glia
Retinal neurons
Retinal pigment cells
Retinal progenitors and Muller glia
Early mesoderm cells
Cardiac mesoderm cells
Cardiomyocytes
Epicardial fat cells
Smooth muscle cells
Vascular endothelial cells
Stromal cells
Metanephric cells
Skeletal muscle cells
Lymphoid cells
Erythroblasts
Megakaryocytes
Myeloid cells
Hepatoblasts
Stellate cells
iPSCs
NES
>= 4
0
<= -4

**Supplementary Figure 7.** GO BP pathway enrichment for each ALK inhibitor from GSEA using post-mash statistics (posterior mean / standard deviation). Each cell of the heatmap shows the Normalized Enrichment Score (NES) for a GO BP gene set for each drug–cell-type combination. Blue indicates upregulation relative to the other four ALK inhibitors; orange indicates downregulation. The top 15 positive and negative NES pathways (i.e. largest max(NES) and most negative min(NES) across cell types with at least one significant cell type in that direction) are shown. Within each up/down block, pathways are ordered by their signed extreme NES. Each row is a cell type and each column is a pathway. * indicates significant enrichment (p-adj < 0.05). Gray boxes indicate cell-type–pathway combinations where NES could not be computed (for example, too few genes overlapped the gene set in that cell type).

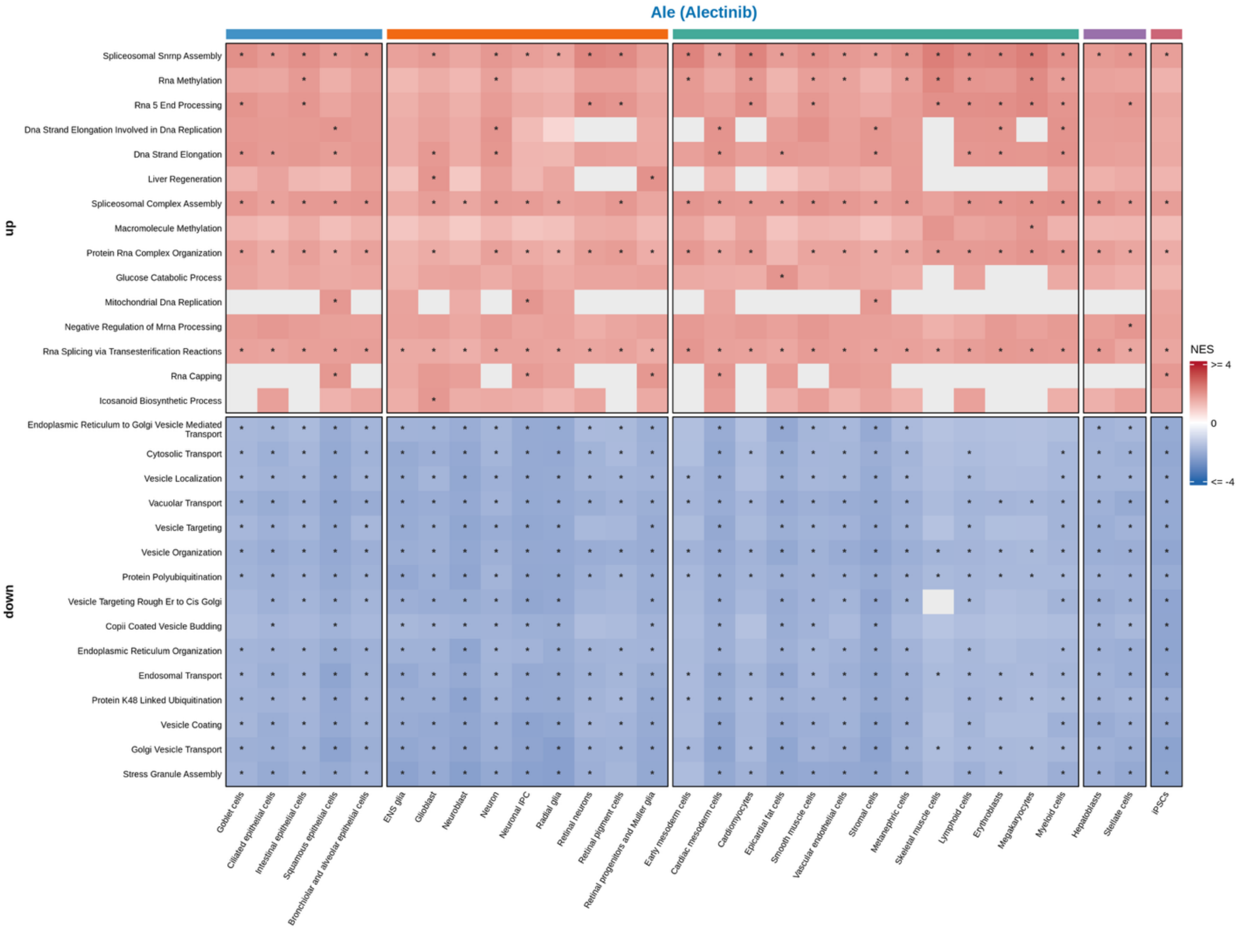

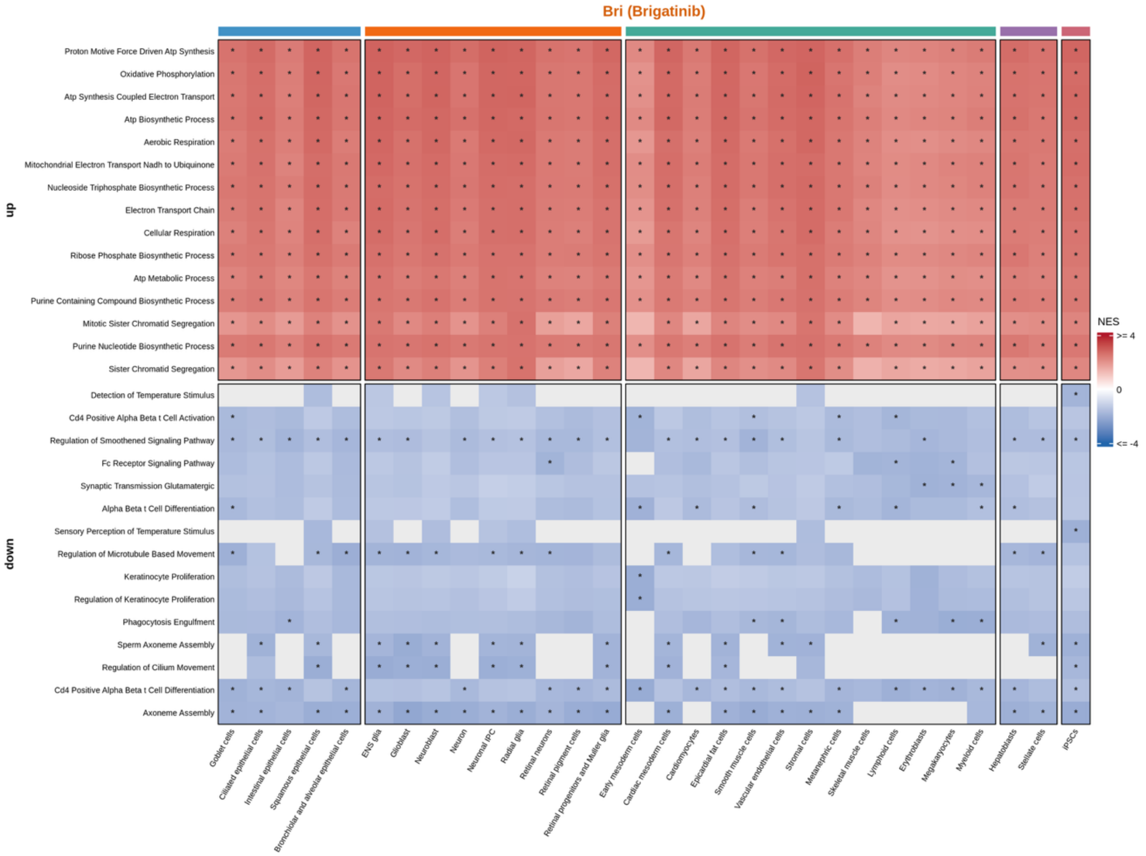

Bri (Brigatinib)
up
down
Proton Motive Force Driven Atp Synthesis
Oxidative Phosphorylation
Atp Synthesis Coupled Electron Transport
Atp Biosynthetic Process
Aerobic Respiration
Mitochondrial Electron Transport Nadh to Ubiquinone
Nucleoside Triphosphate Biosynthetic Process
Electron Transport Chain
Cellular Respiration
Ribose Phosphate Biosynthetic Process
Atp Metabolic Process
Purine Containing Compound Biosynthetic Process
Mitotic Sister Chromatid Segregation
Purine Nucleotide Biosynthetic Process
Sister Chromatid Segregation
Detection of Temperature Stimulus
Cd4 Positive Alpha Beta t Cell Activation
Regulation of Smoothened Signaling Pathway
Fc Receptor Signaling Pathway
Synaptic Transmission Glutamatergic
Alpha Beta t Cell Differentiation
Sensory Perception of Temperature Stimulus
Regulation of Microtubule Based Movement
Keratinocyte Proliferation
Regulation of Keratinocyte Proliferation
Phagocytosis Engulfment
Sperm Axoneme Assembly
Regulation of Cilium Movement
Cd4 Positive Alpha Beta t Cell Differentiation
Axoneme Assembly
Goblet cells
Ciliated epithelial cells
Intestinal epithelial cells
Squamous epithelial cells
Bronchiolar and alveolar epithelial cells
ENS glia
Glioblast
Neuroblast
Neuron
Neuronal IPC
Radial glia
Retinal neurons
Retinal pigment cells
Retinal progenitors and Muller glia
Early mesoderm cells
Cardiac mesoderm cells
Cardiomyocytes
Epicardial fat cells
Smooth muscle cells
Vascular endothelial cells
Stromal cells
Metanephric cells
Skeletal muscle cells
Lymphoid cells
Erythroblasts
Megakaryocytes
Myeloid cells
Hepatoblasts
Stellate cells
iPSCs
NES
>= 4
0
<= -4

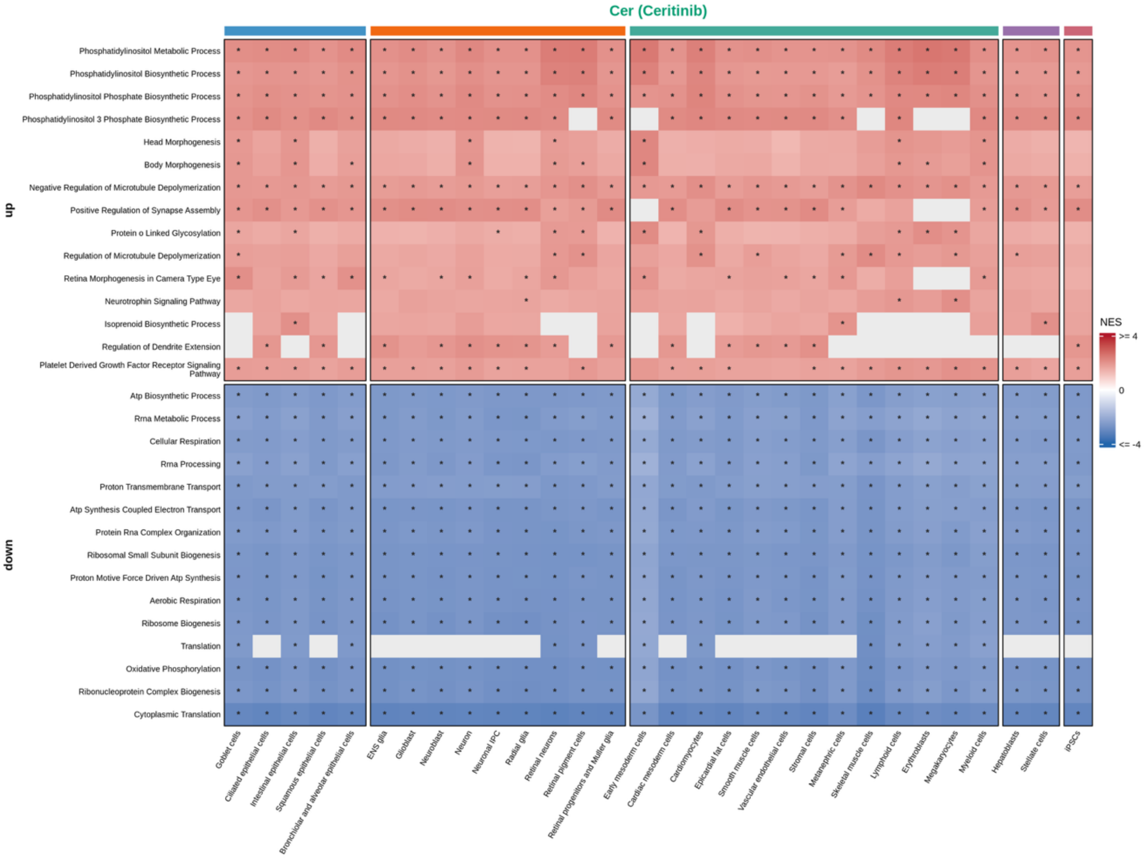
Cer (Ceritinib)
up
down
Phosphatidylinositol Metabolic Process
Phosphatidylinositol Biosynthetic Process
Phosphatidylinositol Phosphate Biosynthetic Process
Phosphatidylinositol 3 Phosphate Biosynthetic Process
Head Morphogenesis
Body Morphogenesis
Negative Regulation of Microtubule Depolymerization
Positive Regulation of Synapse Assembly
Protein o Linked Glycosylation
Regulation of Microtubule Depolymerization
Retina Morphogenesis in Camera Type Eye
Neurotrophin Signaling Pathway
Isoprenoid Biosynthetic Process
Regulation of Dendrite Extension
Platelet Derived Growth Factor Receptor Signaling Pathway
Atp Biosynthetic Process
Rrna Metabolic Process
Cellular Respiration
Rrna Processing
Proton Transmembrane Transport
Atp Synthesis Coupled Electron Transport
Protein Rna Complex Organization
Ribosomal Small Subunit Biogenesis
Proton Motive Force Driven Atp Synthesis
Aerobic Respiration
Ribosome Biogenesis
Translation
Oxidative Phosphorylation
Ribonucleoprotein Complex Biogenesis
Cytoplasmic Translation
NES
>= 4
0
<= -4
Goblet cells
Ciliated epithelial cells
Intestinal epithelial cells
Squamous epithelial cells
Bronchiolar and alveolar epithelial cells
ENS glia
Glioblast
Neuroblast
Neuron
Neuronal IPC
Radial glia
Retinal neurons
Retinal pigment cells
Retinal progenitors and Muller glia
Early mesoderm cells
Cardiac mesoderm cells
Cardiomyocytes
Epicardial fat cells
Smooth muscle cells
Vascular endothelial cells
Stromal cells
Metanephric cells
Skeletal muscle cells
Lymphoid cells
Erythroblasts
Megakaryocytes
Myeloid cells
Hepatoblasts
Stellate cells
iPSCs

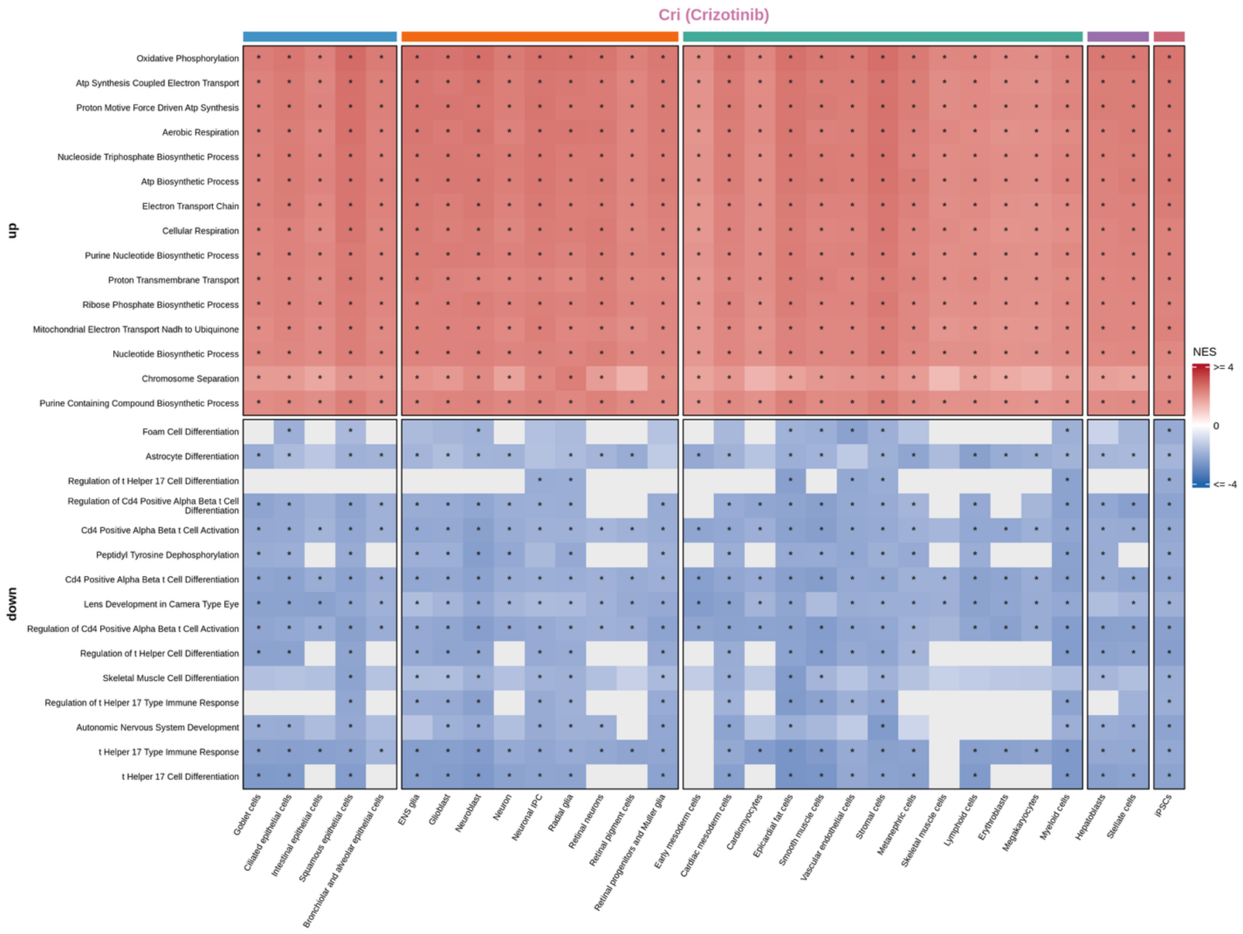

Cri (Crizotinib)
up
down
Oxidative Phosphorylation
Atp Synthesis Coupled Electron Transport
Proton Motive Force Driven Atp Synthesis
Aerobic Respiration
Nucleoside Triphosphate Biosynthetic Process
Atp Biosynthetic Process
Electron Transport Chain
Cellular Respiration
Purine Nucleotide Biosynthetic Process
Proton Transmembrane Transport
Ribose Phosphate Biosynthetic Process
Mitochondrial Electron Transport Nadh to Ubiquinone
Nucleotide Biosynthetic Process
Chromosome Separation
Purine Containing Compound Biosynthetic Process
Foam Cell Differentiation
Astrocyte Differentiation
Regulation of t Helper 17 Cell Differentiation
Regulation of Cd4 Positive Alpha Beta t Cell Differentiation
Cd4 Positive Alpha Beta t Cell Activation
Peptidyl Tyrosine Dephosphorylation
Cd4 Positive Alpha Beta t Cell Differentiation
Lens Development in Camera Type Eye
Regulation of Cd4 Positive Alpha Beta t Cell Activation
Regulation of t Helper Cell Differentiation
Skeletal Muscle Cell Differentiation
Regulation of t Helper 17 Type Immune Response
Autonomic Nervous System Development
t Helper 17 Type Immune Response
t Helper 17 Cell Differentiation
Goblet cells
Ciliated epithelial cells
Intestinal epithelial cells
Squamous epithelial cells
Bronchiolar and alveolar epithelial cells
ENS glia
Glioblast
Neuroblast
Neuron
Neuronal IPC
Radial glia
Retinal neurons
Retinal pigment cells
Retinal progenitors and Muller glia
Early mesoderm cells
Cardiac mesoderm cells
Cardiomyocytes
Epicardial fat cells
Smooth muscle cells
Vascular endothelial cells
Stromal cells
Metanephric cells
Skeletal muscle cells
Lymphoid cells
Erythroblasts
Megakaryocytes
Myeloid cells
Hepatoblasts
Stellate cells
iPSCs
NES
>= 4
0
<= -4

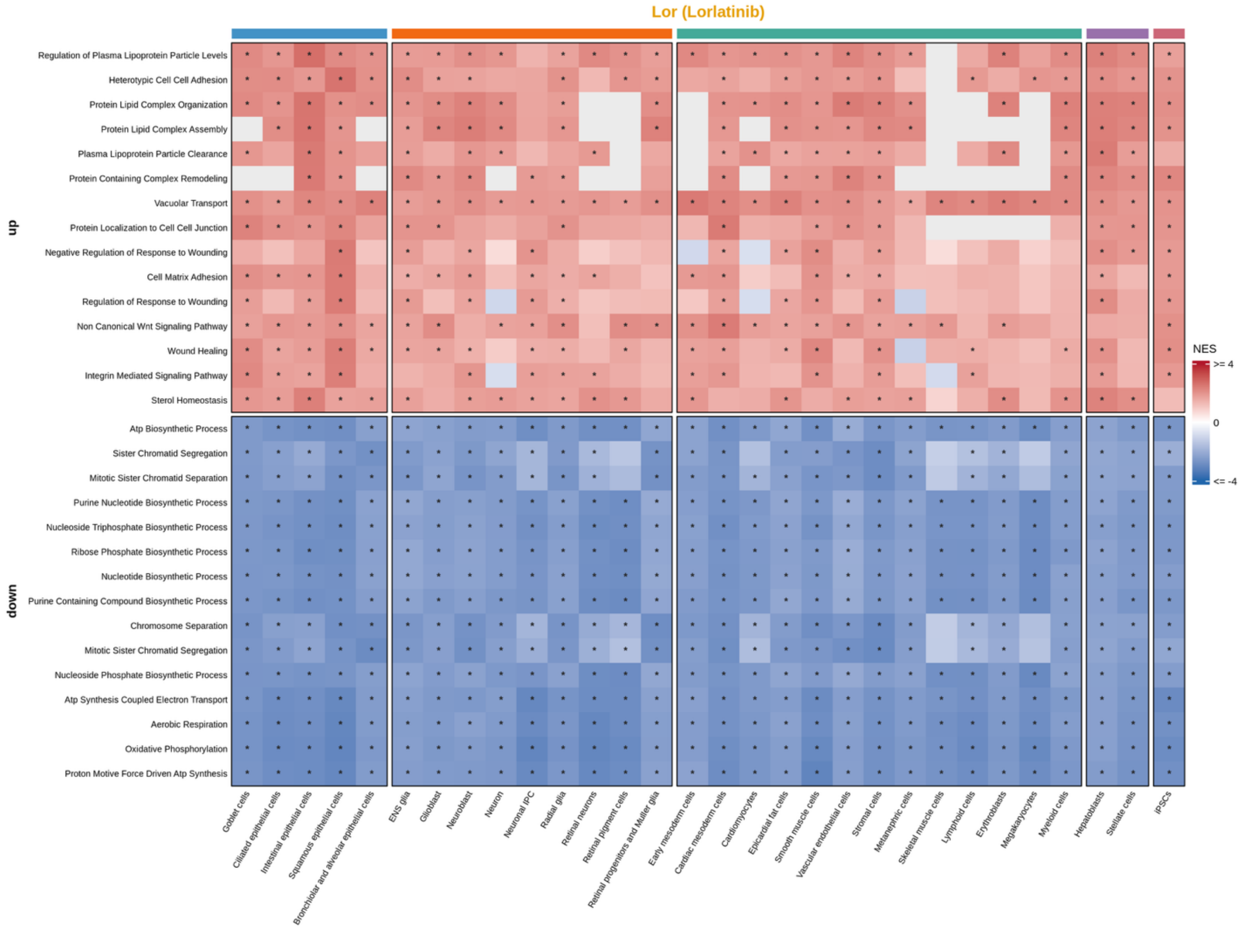

Lor (Lorlatinib)
up
Regulation of Plasma Lipoprotein Particle Levels
Heterotypic Cell Cell Adhesion
Protein Lipid Complex Organization
Protein Lipid Complex Assembly
Plasma Lipoprotein Particle Clearance
Protein Containing Complex Remodeling
Vacuolar Transport
Protein Localization to Cell Cell Junction
Negative Regulation of Response to Wounding
Cell Matrix Adhesion
Regulation of Response to Wounding
Non Canonical Wnt Signaling Pathway
Wound Healing
Integrin Mediated Signaling Pathway
Sterol Homeostasis
down
Atp Biosynthetic Process
Sister Chromatid Segregation
Mitotic Sister Chromatid Separation
Purine Nucleotide Biosynthetic Process
Nucleoside Triphosphate Biosynthetic Process
Ribose Phosphate Biosynthetic Process
Nucleotide Biosynthetic Process
Purine Containing Compound Biosynthetic Process
Chromosome Separation
Mitotic Sister Chromatid Segregation
Nucleoside Phosphate Biosynthetic Process
Atp Synthesis Coupled Electron Transport
Aerobic Respiration
Oxidative Phosphorylation
Proton Motive Force Driven Atp Synthesis
Goblet cells
Ciliated epithelial cells
Intestinal epithelial cells
Squamous epithelial cells
Bronchiolar and alveolar epithelial cells
ENS glia
Glioblast
Neuroblast
Neuron
Neuronal IPC
Radial glia
Retinal neurons
Retinal pigment cells
Retinal progenitors and Muller glia
Early mesoderm cells
Cardiac mesoderm cells
Cardiomyocytes
Epicardial fat cells
Smooth muscle cells
Vascular endothelial cells
Stromal cells
Metanephric cells
Skeletal muscle cells
Lymphoid cells
Erythroblasts
Megakaryocytes
Myeloid cells
Hepatoblasts
Stellate cells
iPSCs
NES
>= 4
0
<= -4

**Supplementary Figure 8**. Number of significant *cis*-eQTLs (lfsr < 0.05) per cell-type–treatment (or control) condition. Darker red indicates a greater number of significant eQTLs. Rows are ordered by mean eQTL count across conditions (highest at top).

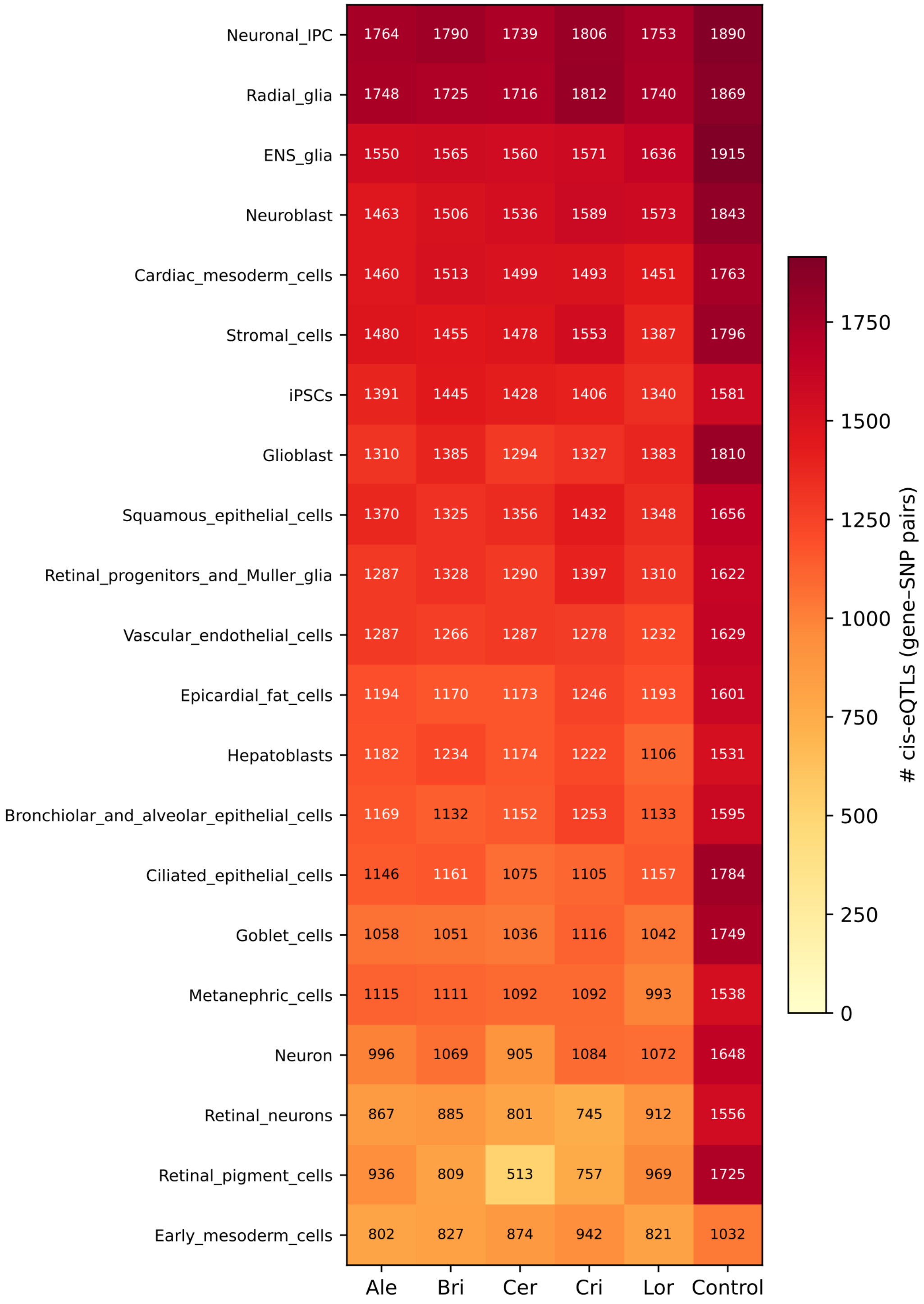

**Supplementary Figure 9**. Standard *cis*-eQTL counts (lfsr < 0.05) per condition plotted against pseudobulk sample size. (A) Treatment conditions: eQTL count vs number of donors contributing to each cell-type–treatment pseudobulk. Each point is one cell-type–treatment combination, colored by drug. (B) Treatment conditions: eQTL count vs total number of cells aggregated into each pseudobulk. (C) Control eQTL counts vs number of donors per cell type; points labeled by cell type. (D) Control eQTL count vs number of cells per cell type. Pearson correlation (r) is shown in the lower right of each panel; gray line and shaded band show the linear fit and 95% confidence interval.

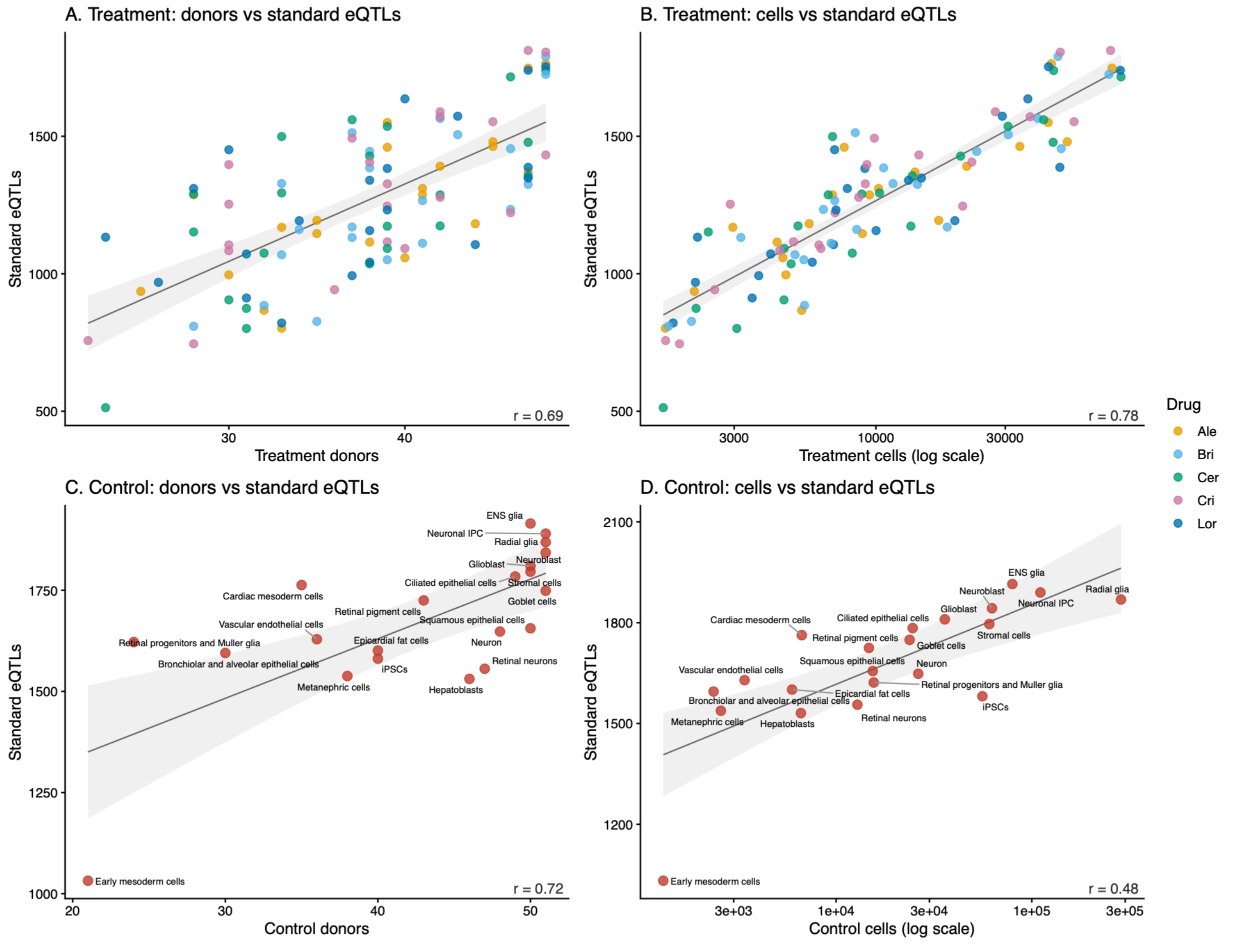

**Supplementary Figure 10.** Concordance between mash-based response eQTL effect sizes and an explicit genotype-by-treatment interaction model. Each panel shows one ALK inhibitor. The x-axis is the difference in mash posterior effect size between treatment and control (Δβ = β_treatment – β_control) and the y-axis is the genotype–treatment coefficient from the interaction model fit in the same cell type. Points are unique gene–SNP pairs classified as response eQTLs for that drug (n = 1,444–1,553 per panel, one point per pair, using the cell type with the lowest lfsr where a pair was a response eQTL in more than one cell type). Dashed line is the ordinary least-squares fit.

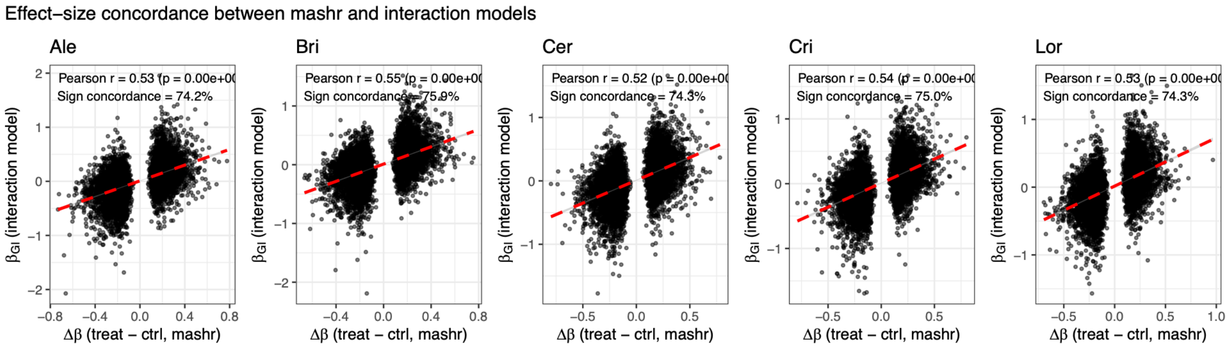

**Supplementary Figure 11**. Density of SNP-to-TSS distance for response and non-response eQTLs for all cell-type–treatment conditions, grouped by germ layer. Distances were calculated within a +/-100kb cis window. Shaded area shows histogram density and lines show density estimates.

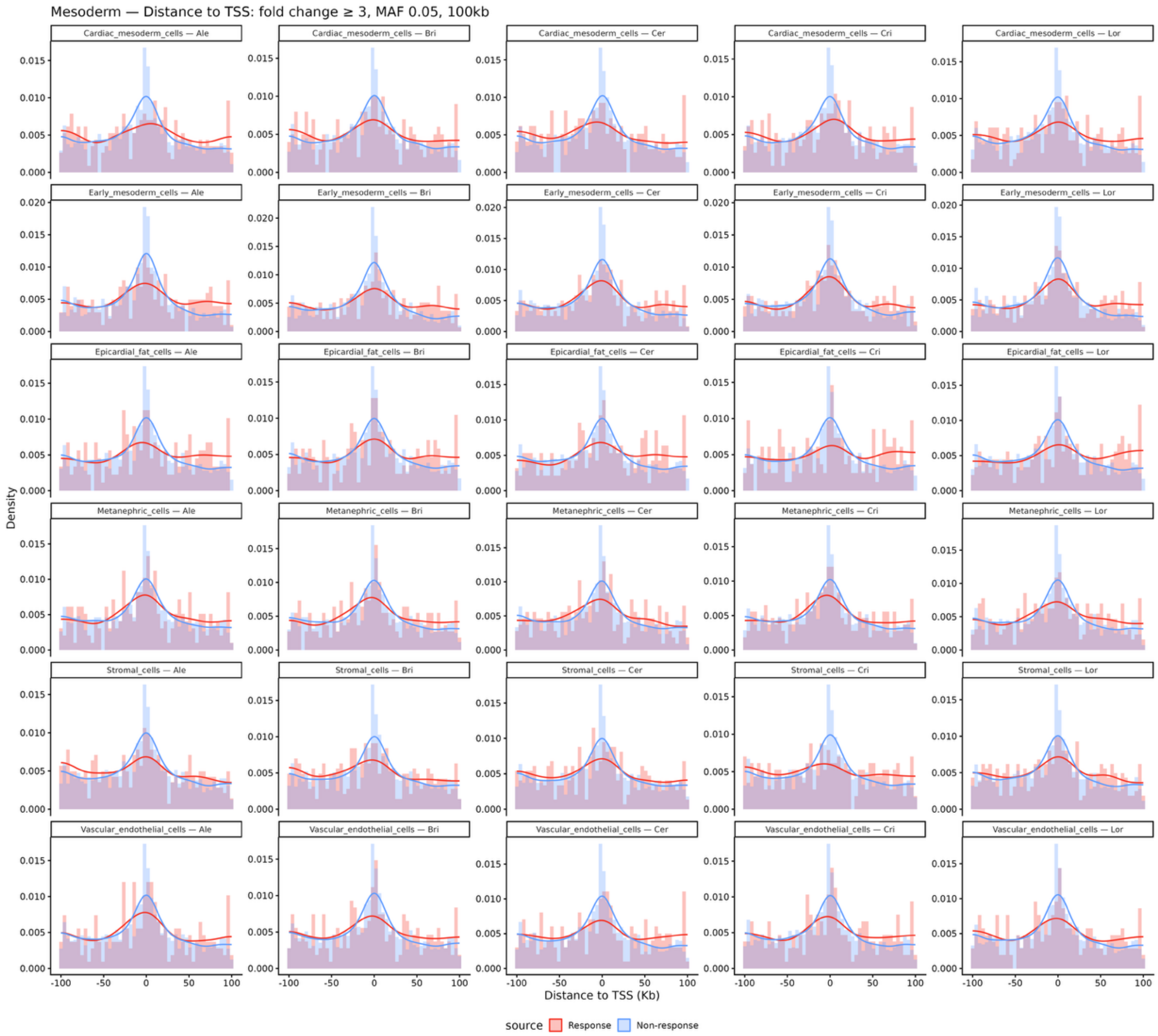

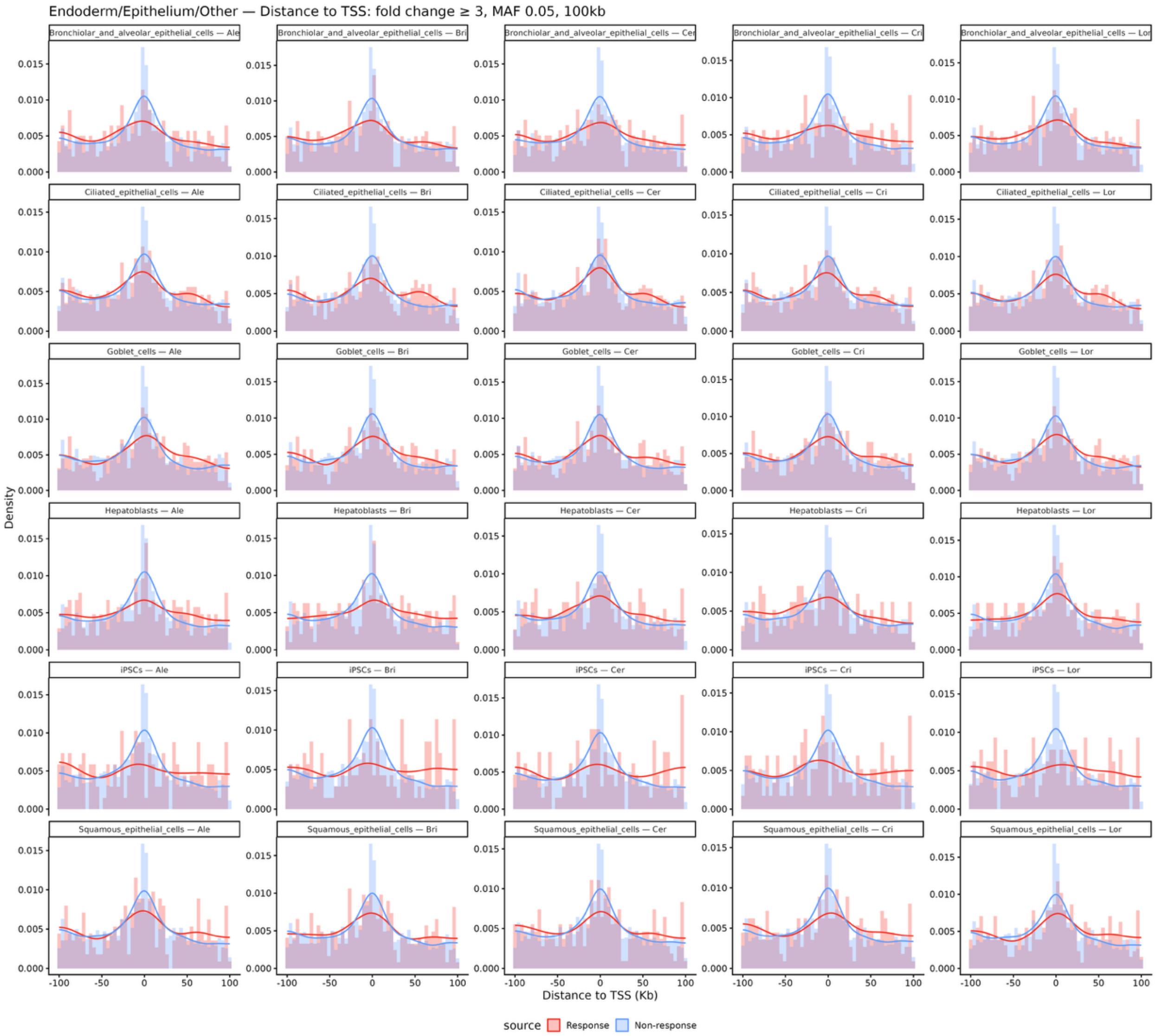
Endoderm/Epithelium/Other — Distance to TSS: fold change ≥ 3, MAF 0.05, 100kb
Bronchiolar_and_alveolar_epithelial_cells — Ale
Bronchiolar_and_alveolar_epithelial_cells — Bri
Bronchiolar_and_alveolar_epithelial_cells — Cer
Bronchiolar_and_alveolar_epithelial_cells — Cri
Bronchiolar_and_alveolar_epithelial_cells — Lor
Ciliated_epithelial_cells — Ale
Ciliated_epithelial_cells — Bri
Ciliated_epithelial_cells — Cer
Ciliated_epithelial_cells — Cri
Ciliated_epithelial_cells — Lor
Goblet_cells — Ale
Goblet_cells — Bri
Goblet_cells — Cer
Goblet_cells — Cri
Goblet_cells — Lor
Hepatoblasts — Ale
Hepatoblasts — Bri
Hepatoblasts — Cer
Hepatoblasts — Cri
Hepatoblasts — Lor
iPSCs — Ale
iPSCs — Bri
iPSCs — Cer
iPSCs — Cri
iPSCs — Lor
Squamous_epithelial_cells — Ale
Squamous_epithelial_cells — Bri
Squamous_epithelial_cells — Cer
Squamous_epithelial_cells — Cri
Squamous_epithelial_cells — Lor
Density
0.000
0.005
0.010
0.015
-100
-50
0
50
100
Distance to TSS (Kb)
source
Response
Non-response

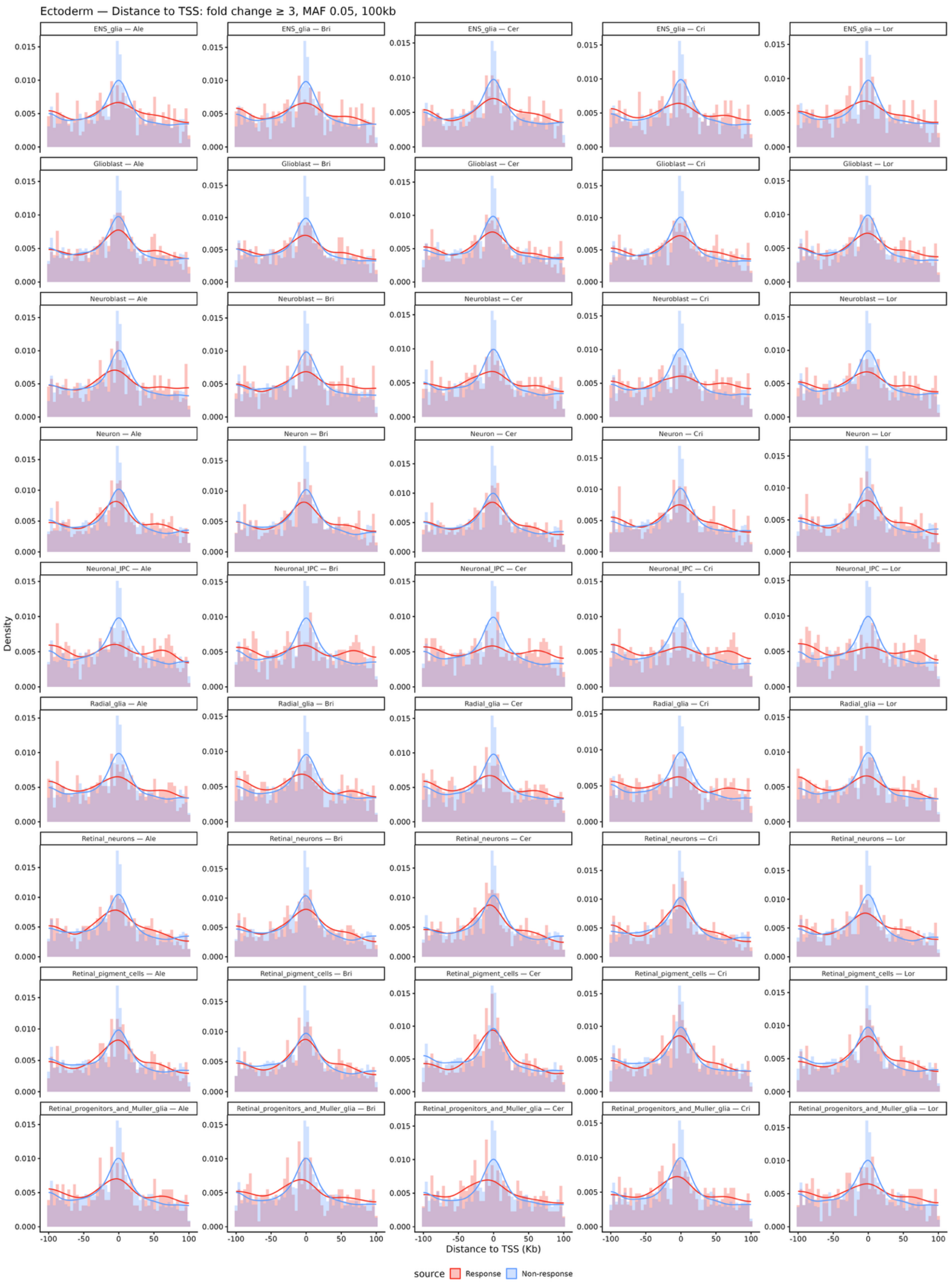
Ectoderm — Distance to TSS: fold change ≥ 3, MAF 0.05, 100kb
ENS_glia — Ale
ENS_glia — Bri
ENS_glia — Cer
ENS_glia — Cri
ENS_glia — Lor
Glioblast — Ale
Glioblast — Bri
Glioblast — Cer
Glioblast — Cri
Glioblast — Lor
Neuroblast — Ale
Neuroblast — Bri
Neuroblast — Cer
Neuroblast — Cri
Neuroblast — Lor
Neuron — Ale
Neuron — Bri
Neuron — Cer
Neuron — Cri
Neuron — Lor
Neuronal_IPC — Ale
Neuronal_IPC — Bri
Neuronal_IPC — Cer
Neuronal_IPC — Cri
Neuronal_IPC — Lor
Radial_glia — Ale
Radial_glia — Bri
Radial_glia — Cer
Radial_glia — Cri
Radial_glia — Lor
Retinal_neurons — Ale
Retinal_neurons — Bri
Retinal_neurons — Cer
Retinal_neurons — Cri
Retinal_neurons — Lor
Retinal_pigment_cells — Ale
Retinal_pigment_cells — Bri
Retinal_pigment_cells — Cer
Retinal_pigment_cells — Cri
Retinal_pigment_cells — Lor
Retinal_progenitors_and_Muller_glia — Ale
Retinal_progenitors_and_Muller_glia — Bri
Retinal_progenitors_and_Muller_glia — Cer
Retinal_progenitors_and_Muller_glia — Cri
Retinal_progenitors_and_Muller_glia — Lor
Density
0.000
0.005
0.010
0.015
-100
-50
0
50
100
Distance to TSS (Kb)
source
Response
Non-response

**Supplementary Figure 12**. Response vs nonresponse cCRE enrichment across three ENCODE candidate cCRE classes. PLS = Promoter-like signature; pELS = proximal enhancer-like signature; dELS = distal enhancer-like signature. Bars show the log2 odds ratio from a two-sided Fisher's exact test comparing the proportion of response versus non-response eQTLs overlapping each cCRE class. Positive values indicate enrichment among response eQTLs; negative values indicate depletion. * = $p < 0.05$; *** = $p < 0.001$

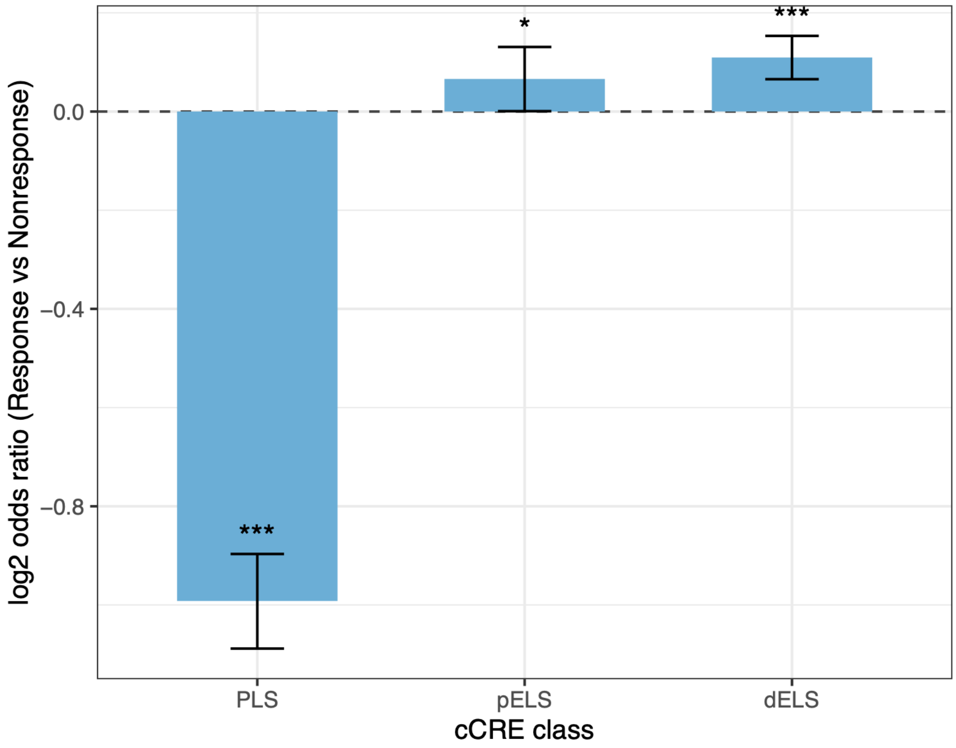
